# Sparse Neuronal C4 Elevation Induces Network-Wide Transcriptomic and Lipid-Droplet Remodeling in the Cortical Microenvironment

**DOI:** 10.1101/2025.05.29.656749

**Authors:** Marta Sánchez-Carbonell, Sonia Bolshakova, Juan E. Ávila-Pagán, Yaseer A. Sabir, Clay Griffin-Derr, Dominik G. Stich, Sofia V. Cieslewicz, Sheyla Esther Carmen Sifuentes, Rhushikesh A. Phadke, Alison Brack, Anna Hasche-Kluender, Paige Caley, Caroline Dias, Alberto Cruz-Martín

**Author notes:** **Corresponding author**: Alberto Cruz-Martín. **These authors contributed equally:** Marta Sánchez-Carbonell,; Sonia Bolshakova,; Juan Ávila-Pagán.

## Abstract

The complement component C4 regulates synaptic refinement and plasticity in the brain and has been implicated in multiple neurological and psychiatric disorders, with especially strong genetic evidence linking elevated *C4A* expression to schizophrenia. However, it remains unclear whether localized C4 elevation in a small neuronal population is sufficient to reshape broader cortical gene-regulatory programs and cellular phenotypes. To address this question, we overexpressed mouse *C4* in approximately 2% of prefrontal cortical neurons using in utero electroporation and performed bulk RNA sequencing on microdissected prefrontal cortex. Despite the sparse manipulation, C4-OE was associated with broad transcriptional remodeling across tissue composed predominantly of untransfected cells, including coordinated changes in cholesterol biosynthesis, axon- and synaptic-associated programs, vascular/ECM pathways, and immune- and stress-related responses. Comparison with a schizophrenia synaptic proteomic dataset identified limited global correspondence but selective gene- and pathway-level overlap in specific molecular programs. Reference-based projection of directional DEG signatures onto a cortical single-cell transcriptomic reference further identified cell-type-associated structure, including prominent astrocyte- and vascular-associated axes and distinct lipid/metabolic, vascular/endothelial–interferon-associated, and immune/stress correlation domains. Spatial validation with MFISH showed increased *Hmgcr* expression in neighboring GFP-negative cells, while PLIN2 imaging revealed increased DAPI-associated/perinuclear lipid-droplet burden within the local C4-OE field. Together, these findings support a model in which sparse neuronal C4 elevation is associated with broader remodeling of the local cortical environment, including a non-cell-autonomous component involving neighboring-cell lipid-associated responses. These results expand the interpretation of C4 risk biology beyond synapse elimination alone and suggest that focal immune-gene dysregulation can engage lipid-homeostatic, stress-related, and cell-type-associated transcriptional programs relevant to cortical circuit vulnerability.

**Author Summary:** Complement component C4 is best known for its role in immune defense, but increasing evidence suggests that altered C4 levels can also influence brain development and disease-relevant neural circuits. Here, we asked what happens when C4 is increased in only a sparse subset of cortical neurons. Using a mouse model with C4 elevation in a small population of prefrontal cortical neurons, we found that this localized perturbation was associated with broad molecular changes across the surrounding cortical field. Transcriptomic analyses revealed coordinated changes in cholesterol biosynthesis and lipid metabolism, synaptic and neuronal programs, vascular-associated pathways, and immune- and stress-related responses. Spatial validation further showed increased *Hmgcr* expression and greater PLIN2-positive lipid-droplet burden in neighboring GFP-negative cells, supporting a non-cell-autonomous component of the response. These findings show that a highly localized increase in one immune-linked gene can ripple across the surrounding cortex, reshaping lipid homeostasis and microenvironmental states linked to circuit vulnerability.

## Introduction

The complement pathway is a key component of the innate immune system, consisting of circulating proteins that, upon activation, trigger a cascade that promotes pathogen clearance and the removal of damaged cells (1,2). Complement proteins play a key role in maintaining immune surveillance within the central nervous system (3). Beyond fighting infections, complement proteins are involved in synapse elimination (4,5), a crucial process for normal brain development and adaptability (6). Microglia, the brain’s resident immune cells, utilize complement proteins such as C1q and C3 to recognize and mark weaker or excess synapses for removal, thereby refining neural circuits during development and in response to experiences (5). In support of this, Stevens et al. (4) showed that knock-out (KO) mice for either C1q, the initiating recognition protein of the classical complement pathway, or the downstream complement protein C3, exhibit deficits in connectivity in the visual thalamus. Furthermore, Schafer et al. (7) demonstrated that microglia actively engulf presynaptic inputs during the peak of retinogeniculate pruning, a process dependent on neural activity and the microglia-specific phagocytic signaling pathway, complement receptor 3 (CR3)/C3.

Sekar et al. (8) showed that structurally diverse *C4* alleles contribute to schizophrenia (SCZ) risk by driving *C4A* expression in the brain, implicating complement-mediated synaptic elimination as a key pathogenic mechanism in this disorder. In support of this, our group and others (9,10) demonstrated that increased levels of the human and mouse homologs of this neuroimmune gene cause decreased connectivity in layer (L) 2/3 pyramidal neurons in the medial prefrontal cortex (mPFC) and deficits in social and cognitive behavior in mice, mirroring human cellular and behavioral pathology (11–16). Importantly, recent findings by Phadke et al. (17) challenge the view that complement acts solely via microglia, showing that *C4* overexpression (C4-OE) impairs synaptic plasticity through an intracellular SNX27-dependent pathway that disrupts AMPAR trafficking and synapse formation. These results suggest that complement may still drive cellular dysfunction and synaptic pathology through previously unrecognized mechanisms.

Global knockout and overexpression models have advanced our understanding of complement-mediated pathology (4,7,18). However, in the peripheral immune system, complement activation often initiates locally, where it can amplify and propagate responses to neighboring cells or tissues (5,19–21). Studying such propagation in the CNS is more challenging not only due to the brain’s complex structure and limited accessibility, but also because it is unclear whether the same mechanisms that govern complement activity in peripheral tissues also govern neural circuits. Given the complexity of neurodevelopmental and neurodegenerative disorders associated with complement pathology (8,11,22–29), model systems in which complement dysregulation can propagate independently of multifactorial disease processes may be critical for disentangling complement-mediated pathology.

To determine whether localized neuronal C4 elevation can reshape broader cortical gene-regulatory programs and cellular phenotypes, we used in utero electroporation (IUE) to overexpress *C4* in approximately 2% of prefrontal cortical neurons, primarily developing layer 2/3 neurons in the medial prefrontal cortex (mPFC), generating a sparse model in which *C4*-overexpressing neurons are surrounded by a majority of untransfected neighboring cells (9,17,30). We then microdissected the electroporated region and performed bulk RNA sequencing (RNA-seq) to identify downstream molecular responses associated with sparse *C4* overexpression (C4-OE).

Despite the limited number of directly transfected neurons, sparse C4-OE was associated with broad transcriptional changes across the local cortical field. These included cholesterol and sterol biosynthesis, synaptic and axonal, vascular/ECM, and directionally distinct immune- and stress-associated transcriptional programs, with cholesterol biosynthesis emerging as one of the most consistent signatures across complementary analyses. Comparison with a SCZ synaptic proteomic dataset revealed limited global molecular correspondence but selective overlap in specific programs, including RNA localization and a subset of Reactome pathways. Reference-based projection of directional DEG signatures onto a cortical single-cell transcriptomic reference further identified neuronal- and non-neuronal-associated transcriptional structure, including prominent astrocyte- and vascular-associated axes. Finally, MFISH and PLIN2 imaging extended these findings beyond bulk RNA-seq by revealing increased *Hmgcr* expression and greater DAPI-associated/perinuclear PLIN2-positive lipid-droplet burden in neighboring GFP-negative cells. Together, these findings support a model in which sparse neuronal C4 elevation is associated with broader transcriptional and lipid-associated remodeling of the surrounding cortical microenvironment.

## Methods

### Ethics statement

All experimental protocols were conducted in accordance with the National Institutes of Health (NIH) guidelines for animal research and were approved by the Boston University (#17–031) and the University of Colorado Anschutz Medical Campus (#01459) Institutional Animal Care and Use Committees.

### Animals

Experimental male and female mice were housed in groups and exposed to a 12-hour light-dark cycle, with the lights turning on at 7 AM and off at 7 PM. Animals had ad libitum access to food and water. The pups remained with their mothers until weaning on Postnatal Day (P) 21. We used wild-type CD-1 mice of both sexes from the CD-1 IGS strain (Charles River; strain code: 022) at P21-23.

### cDNA Constructs

For GFP control, we used a plasmid containing EGFP under the CAG promoter (pCAG-EGFP, Addgene plasmid #11150; http://n2t.net/addgene:11150;RRID:Addgene_11150) (31). The coding sequence for mouse *C4b* (NM_009780.2), synthesized by GenScript, was subcloned into the pCAG expression vector using the InFusion cloning kit (Clontech) to generate the pCAG-mC4 construct (9). The plasmid DNA was purified using the ZymoPure II (Zymo Research) plasmid preparation kit and was resuspended in molecular biology-grade water.

### In Utero Electroporation

In utero electroporations (IUE) targeting L2/3 progenitor cells in the medial prefrontal cortex (mPFC) were performed as previously described (9,17,30). pCAG-EGFP was electroporated to produce the GFP-control condition, and the pCAG-EGFP plasmid was co-electroporated with pCAG-m (mouse) *C4* (*C4b*) to create the C4-OE condition. Using M-FISH, we demonstrated that when we performed IUE with separate plasmids, 99% of the transfected cells expressed mRNAs for all constructs (9,17). This suggests that we can use this gene transfer approach to express multiple constructs in the same cells.

Before the laparotomy, all tools were sterilized using autoclaving. Aseptic techniques were consistently maintained throughout the procedure, and a sterile field was prepared with sterile cloth drapes before surgery began. Timed-pregnant female CD-1 mice at gestation day (E) 16 were anesthetized by inhalation of 4% isoflurane and maintained on 1%–1.5% isoflurane via mask inhalation. The abdomen was sterilized with 10% povidone-iodine and 70% isopropyl alcohol before a vertical incision was made in the skin and then in the abdominal wall. The uterine horn was then exposed, and plasmid DNA was injected into one lateral ventricle by inserting the glass pipette at a 90° angle to the midline of the embryo’s head and injecting 2–4 μl of DNA solution. For GFP-control IUEs, embryos received pCAG-EGFP at 1 μg/μl, corresponding to 2–4 μg of pCAG-EGFP per embryo. For C4-OE IUEs, embryos received pCAG-EGFP at 1 μg/μl together with pCAG-mC4/C4b at 1 μg/μl, corresponding to a GFP:C4 plasmid ratio of 1:1 and 2–4 μg of each plasmid per embryo. Next, four square pulses (pulse duration: 50 ms, pulse amplitude: 36 V, interpulse interval: 500 ms) were delivered to the head of the embryo using a custom-built tripolar electroporator (30,32). The embryos were regularly moistened with warm sterile PBS during the surgical procedure. After electroporation, the embryos and uterine horn were gently placed back in the dam’s abdominal cavity, and the muscle and skin were sutured using absorbable and nonabsorbable sutures, respectively. Finally, the dams were allowed to recover in a warm chamber for 1 hour, then returned to their cages.

### In Utero Electroporation Efficiency

To estimate the proportion of neurons transfected via IUE, we quantified GFP-positive cells within defined regions of interest (ROIs) in the PFC at P21 (9). We analyzed six model brain sections (9), each 45 μm thick, which contained a total of 108 ROIs measuring 1 mm by 0.5 mm and tiling the prefrontal cortex. The total sampled tissue volume was approximately 2.43 mm³. Within this volume, we identified an average of about 4,000 GFP-positive neurons (9). Based on a reported mouse cortical neuronal density of approximately 92,616 ± 25,000 neurons/mm³ (33), we estimate that the sampled region contained approximately 164,000–286,000 neurons. This results in an average transfection rate of around 1.78%, with a range of roughly 1.40% to 2.43%. These findings confirm the expected sparse labeling achieved through IUE, enabling the targeted manipulation of a small subset of neurons for subsequent molecular and circuit-level analyses.

For MFISH experiments, local transfection density was also quantified within the GFP-positive hotspot selected for spatial analysis. GFP-positive C4-OE cells were counted within the analyzed region of interest, and the total sampled cell population was estimated from DAPI-positive nuclei in the same region. Local transfection density was calculated as the percentage of GFP-positive cells among all DAPI-positive cells within the sampled hotspot. Using this approach, GFP-positive cells represented approximately 7% of the locally sampled cell population. This hotspot-specific estimate is expected to be higher than the broader estimate of approximately 2% electroporation efficiency because the larger PFC sampling strategy included the GFP-positive hotspot, peri-transfected regions, and more sparsely labeled portions of the mPFC.

### Tissue Extraction for Bulk RNA Sequencing

P21-23 brains were rapidly extracted following transcardiac perfusion using ice-cold PBS. NMDG slicing solution consisted of 92 mM NMDG, 2.5 mM KCl, 1.25 mM NaH_2_PO_4_, 30 mM NaHCO_3_, 20 mM HEPES, 25 mM glucose, 2 mM thiourea, 5 mM Na-ascorbate, 3 mM Na pyruvate, 0.5 mM CaCl_2_·2H2O, and 10 mM MgSO_4_·7H2O. Frontal sections (200 μm), primarily including the prelimbic, infralimbic, anterior cingulate cortex, and supplementary motor cortex, were cut on a vibratome (Leica VT1000 S) and kept in NMDG slicing solution bubbled with 95% O_2_/5% CO_2_ (295–305 mOsm). Sections were trimmed under a wide-field microscope to isolate the transfected region (containing GFP-positive cells) of the tissue. Next, the GFP-positive tissue was immediately flash-frozen on dry ice.

### Bulk RNA-seq Analysis

Aligned BAM files were quantified using featureCounts (v2.0.1) (34), with the Gencode vM23 annotation GTF file used as the reference for gene identification and read assignment. Ensembl gene identifiers were converted to MGI gene symbols using the org.Mm.eg.db annotation package. When multiple Ensembl identifiers mapped to the same gene symbol, counts were summed across identifiers prior to downstream analysis. Genes with zero variance across samples and genes with nonzero counts in one or fewer samples were excluded before differential-expression analysis. Exploratory quality-control analyses included assessment of library size and the relationship between gene-level mean expression and variance.

Downstream analyses were performed in R (v4.2.1; R Foundation for Statistical Computing). Differential expression was analyzed using DESeq2 (35), with sex included as an additive covariate in the design (∼ sex + genotype). Control was specified as the reference genotype. The Control group consisted of four mice (2 females and 2 males), whereas the C4-OE group consisted of four male mice. DESeq2 was run using the standard DESeq() workflow, and genotype effects were extracted as the C4-OE versus Control coefficient using results(). No log₂ fold-change shrinkage was applied; therefore, reported log₂ fold changes correspond to the estimates returned directly by DESeq2. Differentially expressed genes were defined using a Benjamini–Hochberg-adjusted P value < 0.05 without an additional fold-change threshold. DESeq2 was selected because it provides a well-established framework for count-based RNA-seq differential-expression analysis and has been widely applied in neurodevelopmental transcriptomic studies (35,36).

Sample sex was inferred from raw RNA-seq counts before downstream filtering using the reciprocal expression patterns of *Xist* (37) and the Y-linked gene *Ddx3y* (38). Samples with *Xist* counts >1,000 and *Ddx3y* counts <100 were classified as female, whereas samples with *Ddx3y* counts >1,000 and *Xist* counts <100 were classified as male. These inferred sex assignments were recorded in the sample metadata used for downstream analysis and incorporated into the DESeq2 design as the sex covariate.

Because all C4-OE samples were male, whereas Control samples included both males and females, sex and genotype were partially confounded in the experimental design. Inclusion of sex as an additive covariate allowed the genotype coefficient to be estimated after accounting for the sex effect evident within the Control group; however, this design cannot fully distinguish a genotype-associated effect from an effect that is specific to males. This limitation is considered separately in the interpretation of the results.

To identify genes for which genotype-associated differences might also be influenced by sex, we performed a secondary DESeq2 analysis restricted to the four Control samples using a ∼ sex design. Genes with an adjusted P value < 0.05 in the male-versus-female Control comparison were flagged as potentially sex-confounded in the primary DESeq2 results table. Because this secondary comparison included only two males and two females, the flag was used as a screening criterion rather than as a confirmatory test of sex-dependent expression.

Variance-stabilizing transformation was performed using DESeq2 with ‘blind = FALS’, and the resulting expression matrix was used for principal component analysis. For visualization of the most significant differential-expression patterns, the 50 genes with the smallest adjusted P values were selected.

Normalized counts were log₂-transformed after addition of a pseudocount of 1 and standardized within each gene before hierarchical heatmap visualization.

Predefined sex-linked genes were retained and flagged in the complete DESeq2 results table for transparency rather than being removed from the primary differential-expression analysis. These genes were excluded from the primary downstream GSEA ranked list and from the significant-gene input lists used for GO over-representation analysis.

### Gene Set Enrichment Analysis

Gene set enrichment analysis (GSEA) was performed using the fgsea package (v1.22.0) (39), implementing the GSEA framework originally described by Subramanian et al. (40). Genes were ranked by the DESeq2 Wald statistic (stat), defined as the estimated log₂ fold change relative to its standard error, rather than by raw log₂ fold change. This ranking strategy incorporates both the direction and uncertainty of the differential-expression estimate and reduces the influence of low-count, high-variance genes at the extremes of the ranked list. Genes with NA adjusted P values following DESeq2 testing were excluded from the ranked list, as were predefined sex-linked genes.

GSEA was performed using Mouse MSigDB v2023.2 gene-set collections (41), including Hallmark gene sets (42), canonical pathway, Gene Ontology biological process, regulatory-target, and cell-type signature collections. The analyses presented here focus primarily on the Hallmark, canonical pathway, and Gene Ontology biological process collections. Gene sets containing fewer than 15 or more than 500 genes were excluded. Gene sets with a Benjamini–Hochberg-adjusted P value < 0.05 were considered significantly enriched.

### Gene Ontology Over-Representation Analysis

Gene Ontology over-representation analysis (ORA) was performed using clusterProfiler (43). Significant DEGs were defined by Benjamini–Hochberg-adjusted P < 0.05 without an additional absolute log₂ fold-change threshold. No additional fold-change cutoff was applied because the analysis was intended to capture coordinated biological programs that may involve modest but consistent expression changes across multiple genes.

ORA was performed separately on the combined significant DEG set, the upregulated DEG set, and the downregulated DEG set because biological processes associated with the two directions of transcriptional change can differ substantially. Predefined sex-linked genes, predicted genes with symbols beginning with Gm, and genes with symbols ending in Rik were excluded from the significant-gene input lists. These exclusions were not applied to the background universe.

For all ORA analyses, the background universe consisted of all genes that survived preprocessing and were actually tested by DESeq2, rather than all genes represented in the annotation database. This approach restricts the statistical background to genes that had the opportunity to be identified as differentially expressed in the present RNA-seq experiment and avoids enrichment biases associated with using the entire annotated genome as the reference population.

Biological process, cellular component, and molecular function ontologies were evaluated using org.Mm.eg.db, with Benjamini–Hochberg-adjusted P < 0.05 defining significant enrichment. For focused analysis of synaptic biological processes, synaptic GO terms were defined using prespecified parent GO categories together with all of their descendant terms in the GO hierarchy, rather than by keyword matching of term names.

Because ORA evaluates a significance-thresholded subset of genes whereas GSEA evaluates coordinated changes across the full ranked gene list, agreement between the two approaches was interpreted as evidence that an enrichment signal was not dependent on the DEG significance cutoff, rather than as independent statistical replication.

### Weighted Gene Co-expression Network Analysis

Weighted gene co-expression network analysis (WGCNA) was performed using the WGCNA R package (version 1.74) (44) on the variance-stabilized expression matrix described above. Before network construction, genes with a mean DESeq2-normalized count <20 across samples were excluded to reduce the contribution of low-expression genes, leaving 15,729 genes for network construction. This filtering was based only on expression level and was independent of genotype and differential-expression significance. Genes and samples were subsequently assessed using goodSamplesGenes() before network construction.

Soft-thresholding behavior was evaluated across powers 1–20 using signed scale-free topology fit and mean connectivity. Signed scale-free topology fit did not reach the conventional threshold of 0.90 at any power examined (maximum 0.875), so criterion-based selection of a single power was not possible. Networks were therefore independently reconstructed across powers 12– 18 while holding the input gene set, minimum module size, and merge cut height constant, and the resulting partitions were compared using the adjusted Rand index and a size-weighted best-match Jaccard similarity.

A soft-thresholding power of 15 was selected for the primary network analysis because it achieved a signed scale-free topology fit of 0.833 and produced the most representative partition among the solutions examined, showing the highest mean adjusted Rand index relative to the alternative powers (0.356, versus 0.289–0.339 for the other powers). Module-specific stability at power 15 was quantified using the mean best-match Jaccard similarity of each module across the alternative network solutions. These sensitivity analyses were used to evaluate relative module stability and supported power 15 as a representative network solution rather than identifying a uniquely optimal soft-thresholding power.

Co-expression modules were identified using blockwiseModules() with a signed network, a single block encompassing all input genes, deepSplit = 2, pamRespectsDendro = FALSE, reassignThreshold = 0, a minimum module size of 200 genes, and a module-merging cut height of 0.25. Genes not assigned to a module were designated as grey and excluded from module-level biological interpretation.

Module eigengenes were calculated using moduleEigengenes() and standardized across samples as Z scores for visualization. Module–trait relationships were evaluated by correlating module eigengenes with genotype and sex. Pearson correlations and corresponding P values were calculated using WGCNA, and genotype-associated P values were adjusted across modules using the Benjamini–Hochberg method. These associations were interpreted in the context of the partial sex–genotype confounding described above.

GO Biological Process enrichment was performed separately for each non-grey module using clusterProfiler. Genes assigned to each module were used as the query set, with the 15,696 genes returning a non-missing DESeq2 adjusted P value used as the background universe. This background was nearly identical to the 15,729 genes retained for WGCNA network construction; the 33-gene difference reflected genes retained for network construction but removed by DESeq2 independent filtering. Benjamini–Hochberg-adjusted P < 0.05 defined significant enrichment. Significant module-level GO terms were combined for functional annotation and visualization.

### Cross-species comparison of the C4-OE transcriptomic signature with the schizophrenia synaptic proteome

To assess whether molecular programs altered by sparse neuronal C4 elevation overlap with those detected at synapses in SCZ, we compared the mouse C4-OE cortical transcriptome with the synaptic proteomic dataset reported by Aryal et al. (45). Protein-level data were obtained from Table S2 (45) and standardized to one representative protein entry per gene. For genes represented by multiple protein entries, the representative entry was selected based on measurement support rather than statistical significance, prioritizing spectral count, followed by number of unique peptides, sequence coverage, and accession number as a deterministic final tie-breaker.

The mouse input consisted of the sex-adjusted DESeq2 results described above. One record was retained per mouse gene, with duplicate gene-symbol entries, when present, resolved by retaining the row with the highest mean normalized expression (baseMean), independently of statistical significance.

Human–mouse orthologs were mapped using Ensembl BioMart. The analysis was restricted to high-confidence one-to-one orthologs with orthology confidence equal to 1 and unique mappings in both directions. Ensembl release 116 was used, and the resulting ortholog table was frozen for reproducibility. The shared measurable universe comprised ortholog pairs for which the protein was measured in the SCZ synaptic proteomic dataset and the corresponding mouse gene was present in the C4-OE DESeq2 analysis, yielding 7,498 matched genes.

For pathway-level comparisons, genes in the SCZ synaptic proteome were ranked by the SCZ-versus-control proteomic t statistic, whereas mouse genes were ranked by the DESeq2 Wald statistic. Mouse genes were represented by their corresponding human ortholog symbols so that both datasets could be evaluated using identical human gene sets.

Preranked gene set enrichment analysis was performed using fgseaMultilevel with human MSigDB Gene Ontology Biological Process gene sets obtained through msigdbr. Gene sets were restricted to the shared measurable universe, and pathways containing 15–500 measurable genes were tested. Separate enrichment analyses were performed for the SCZ synaptic proteome and mouse C4-OE transcriptome using identical pathway definitions and the same shared measurable universe. Statistical significance was defined as Benjamini–Hochberg-adjusted P < 0.05.

Global pathway correspondence was evaluated across all GO Biological Process pathways tested in both datasets, irrespective of statistical significance. Spearman correlation between normalized enrichment scores (NES) was used as the primary measure of global correspondence, with Pearson correlation examined secondarily. Exact GO pathways were additionally classified according to whether they reached FDR < 0.05 in both datasets and whether their NES values had the same or opposite direction.

For leading-edge comparison of the RNA localization pathway, leading-edge genes were extracted independently from the SCZ and mouse GSEA results, and the shared leading-edge core was defined as the intersection of the two gene sets. The number of shared leading-edge genes and their Jaccard overlap were calculated, and the corresponding gene-level proteomic and transcriptomic statistics were retained for comparison.

To assess whether cross-species correspondence extended beyond GO ontology structure, the same ranked gene lists were also analyzed using MSigDB Hallmark and Reactome gene sets. Hallmark pathways containing 10–500 measurable genes and Reactome pathways containing 15– 500 measurable genes were tested using fgseaMultilevel. Exact pathway matches were identified across datasets, and pathways significant at FDR < 0.05 in both were retained for comparison. Shared significant Reactome pathways were further grouped into broader biological categories based on pathway annotation, and shared leading-edge genes were calculated for each pathway.

As a complementary gene-level analysis, overlap between significantly altered proteins and transcripts was tested within the same shared ortholog universe. The primary comparison defined SCZ-associated differentially expressed proteins at FDR < 0.10 and mouse C4-OE differentially expressed genes at adjusted P < 0.05; a stricter SCZ FDR < 0.05 threshold was examined as a sensitivity analysis. Enrichment of overlapping genes was tested using a one-sided Fisher exact test. Directional concordance among overlapping genes was summarized descriptively based on the sign of the proteomic and transcriptomic effect estimates.

### Cell-type-aware projection of bulk RNA-seq signatures using single-cell reference data

To contextualize the bulk RNA-seq changes observed following sparse C4-OE within known cortical cell-type transcriptional programs, we adapted a previously published STAR Protocol for cell-type-aware projection of bulk RNA-seq gene signatures using single-cell reference data (46). This approach leverages publicly available scRNA-seq datasets to map bulk DEGs onto established cell-type-associated transcriptional landscapes, thereby providing a framework for identifying cortical cell classes and gene modules represented in the C4-OE transcriptional response.

### Construction of a Cortical Single-Cell Reference Dataset

Single-cell transcriptomic reference data were obtained from the Allen Brain Map Mouse Whole Cortex and Hippocampus SMART-seq dataset (2019) with the 10x–SMART-seq taxonomy (2020) (47). Raw gene expression matrices (matrix.csv) and associated cell metadata (metadata.csv) were downloaded from the Allen Brain Map portal and imported into R (v4.1.1). Gene expression counts were converted into a Seurat object (Seurat v4.0.4) without initial filtering to preserve all annotated cell populations (48,49).

Because bulk RNA-seq samples were derived exclusively from microdissected PFC tissue, the reference single-cell RNA-seq dataset was curated to remove hippocampal neuronal populations and retain cortical and broadly distributed non-neuronal cell types. Reference regions were expanded to include the broader cortex rather than being restricted to the PFC alone, thereby increasing representation of cortical cell classes while preserving cell-type identity defined by transcriptomic subclass rather than cortical areal origin. This strategy provided broader representation of cortical cell classes for projection of PFC-derived gene signatures while maintaining transcriptomic cell-class identity. Specifically, cells were filtered based on region_label and subclass_label metadata to exclude hippocampal excitatory neuron subclasses, including CA1, CA2, CA3, and dentate gyrus populations.

Following filtering, the retained cells comprised both neuronal and non-neuronal populations, including astrocytes, oligodendrocytes, microglia, endothelial cells, mural cells (VLMC), cortical interneurons, and pyramidal neurons. Subclass labels without associated cells were removed, and remaining subclasses were consolidated into 19 transcriptomic subclass groups: non-neuronal populations (Astrocytes, Endothelial, Microglia, Oligodendrocytes, and Mural [combining VLMC and SMC-Peri]); inhibitory interneuron populations (Lamp5, Pvalb, Sncg, Sst, Vip, Sst_Chodl, and Other GABA [Meis2]); and excitatory neuronal populations (L2/3_IT, L4/5_IT, L5_PT, L5/6_NP, L6_CT, L6_IT, and Cajal-Retzius). Cell identities were reassigned accordingly and reordered alphabetically for consistency.

Gene expression values were normalized using log normalization with a scale factor of 10,000, and the top 2,000 variable genes were identified using Seurat’s vst feature-selection method. To confirm successful sub-class labeling, a dot plot was generated using Seurat’s DotPlot function with canonical marker genes as features. Data were scaled and centered prior to dimensionality reduction. PCA was performed on variable genes using Seurat’s RunPCA function with a fixed random seed for reproducibility. An elbow plot of variance explained across 50 principal components was inspected, and 14 PCs were retained for downstream nonlinear dimensionality reduction based on the inflection point of the elbow plot.

Nonlinear dimensionality reduction was subsequently performed using t-SNE and UMAP based on the first 14 PCs, with fixed random seeds used for reproducibility. Dimensionality reduction plots (PCA, tSNE, and UMAP) were generated using Seurat’s DimPlot function and combined into a single composite figure using the cowplot and patchwork packages. Distinct colors were assigned to each major cell-type category to facilitate visualization. The resulting figure illustrates the separation and clustering of major cortical cell types within the curated single-cell reference dataset, providing a framework for mapping bulk RNA-seq gene signatures onto cell-type-associated expression profiles.

### Projection of directional C4-OE bulk RNA-seq signatures onto the cortical single-cell reference

To determine whether C4-OE-regulated bulk RNA-seq signatures mapped onto cortical cell-type-associated transcriptional structure, we projected directional DEG signatures onto the curated cortical single-cell reference described above. This analysis was designed to identify reference-cell transcriptional structure represented within the bulk RNA-seq signatures and was not used to infer changes in cell-type abundance.

Bulk RNA-seq differential expression results were stratified into two directional signatures based on the sign of the log2 fold change. Genes expressed at lower levels in C4-OE tissue relative to GFP control tissue were defined as the C4-OE-downregulated signature, whereas genes expressed at higher levels in C4-OE tissue relative to GFP control tissue were defined as the C4-OE-upregulated signature. Gene symbols from each directional signature were intersected with genes present in the curated cortical reference, ranked by log2 fold change within each direction, and the 250 genes with the largest directional effect sizes were retained as the input signatures for downstream projection and correlation analyses. This fixed gene-list size was used to reduce bias introduced by unequal directional DEG-list lengths.

For each directional DEG signature, normalized expression values for the selected genes were extracted from the curated cortical reference. Expression values were scaled across reference cells using Seurat’s ScaleData function, and principal component analysis was performed separately for the C4-OE-downregulated and C4-OE-upregulated signatures. PCA embeddings were visualized using Seurat’s DimPlot, with individual reference cells colored by the consolidated cortical cell-class annotations defined above. These DEG-defined PCA spaces were used to visualize how each directional bulk RNA-seq signature mapped onto cortical reference-cell transcriptional profiles.

To summarize the distribution of cortical reference cell classes across each DEG-defined PCA space, we performed PC coordinate-binning analysis. For each directional projection, reference cells were binned independently along PC1 and PC2 using four equal-width coordinate intervals corresponding to the lowest, low-mid, high-mid, and highest coordinate ranges for each principal component. Within each coordinate bin, cells were grouped by consolidated cortical cell class. The number of cells assigned to each class was counted, and percentages were calculated relative to the total number of reference cells in that bin. Cell-type percentages across PC coordinate bins were visualized as stacked bar plots. This analysis provided a descriptive summary of cell-type-associated structure across the DEG-defined PCA spaces and was not interpreted as evidence for altered cell-type abundance in the bulk RNA-seq samples.

### Correlation-domain analysis of C4-OE-regulated gene signatures

To visualize gene–gene correlation structure within each directional C4-OE DEG signature, expression values for the selected genes were extracted from the normalized data layer of the curated cortical Seurat reference object. Genes with zero variance across reference cells were excluded prior to correlation analysis. Pairwise Pearson correlation coefficients were then calculated among all valid genes across reference cells. For each directional signature, the resulting gene–gene correlation matrix was converted into a distance matrix using Pearson correlation distance, defined as 1 − r, and hierarchically clustered using average linkage clustering.

Correlation-domain assignments were generated from the hierarchical clustering structure using dynamic tree cutting. Dendrogram clusters were identified using cutreeDynamic, with a minimum cluster size of five genes. For the final displayed heatmaps, the C4-OE-downregulated correlation matrix was cut using deepSplit = 2, whereas the C4-OE-upregulated correlation matrix was cut using deepSplit = 1. Gene-domain assignments were used to annotate rows and columns of the corresponding heatmaps.

Correlation heatmaps were plotted using pheatmap, with rows and columns ordered by the same hierarchical clustering structure. Domain annotations were shown for both rows and columns. Color breaks were constructed to place zero at the transition between negative and positive correlations and to span the observed Pearson correlation range for each directional matrix. The resulting C4-OE-downregulated and C4-OE-upregulated correlation heatmaps were used to visualize directional gene–gene correlation structure.

Selected correlation domains were annotated on the heatmaps based on clustering structure, gene content, and biological interpretability. These annotations were used to summarize prominent correlation structure within each directional DEG signature rather than to define discrete cell populations.

### Multiplex fluorescence in situ hybridization and single-cell transcript quantification Tissue preparation and sectioning

Brains were collected after transcardiac perfusion with phosphate-buffered saline (PBS), followed by perfusion with 4% paraformaldehyde (PFA) in PBS. Brains were postfixed in 4% PFA for 24 h at 4°C, cryoprotected in 30% sucrose until they sank, embedded in O.C.T. compound, flash-frozen using ethanol and dry ice, and stored at −80°C. Before sectioning, tissue was equilibrated at −20°C for 30 min. Coronal sections were cut at 15 μm using a Leica CM1950 cryostat at −20°C, wet-mounted in RNase-free PBS onto Fisherbrand Superfrost Plus microscope slides, and stored at −80°C until use.

### Multiplex RNAscope staining

Multiplex fluorescence in situ hybridization was performed using the RNAscope Multiplex Fluorescent V2 assay (Advanced Cell Diagnostics, ACD) according to the manufacturer’s instructions and optimized laboratory protocols. Sections were equilibrated to room temperature for 10 min and baked for 30 min at 60°C. After a 5-min PBS wash, sections were postfixed in 4% PFA for 15 min at 4°C, followed by graded ethanol dehydration in 50%, 70%, and 100% ethanol. Slides were treated with hydrogen peroxide for 10 min at room temperature, subjected to target retrieval in a steamer at 99°C for 5 min, briefly submerged in 100% ethanol, and allowed to air dry overnight.

After an additional 30-min baking step at 60°C, sections were treated with Protease Plus for 15 min at 40°C. The following probes were hybridized for 2 h at 40°C: *Hmgcr* (Mm-Hmgcr, 518921), *Egfp* reporter transcript (EGFP-O4-C3, 538851-C3), and *Fam107a* (Mm-Fam107a-C4, 519011-C4). Signal amplification was performed using AMP 1–3 and HRP reagents. Fluorescent signals were developed using TSA-based Opal fluorophores: Opal 570 for *Hmgcr*, Opal 520 for *Egfp*, and Opal 780 for *Fam107a*. Each fluorophore development step was followed by HRP blocking for 15 min at 40°C. Sections were counterstained with DAPI for 30 s and mounted using ProLong Glass Antifade Mountant.

### Imaging and image analysis

High-resolution imaging was performed using an Olympus VS200 high-throughput slide scanning microscope equipped with a 60× oil-immersion objective, NA 1.42. Fluorescence was captured using a Hamamatsu ORCA-Fusion Gen III sCMOS detector with the following emission filters: DAPI, 413–451 nm; FITC, 497–532 nm for Opal 520; TRITC, 577–613 nm for Opal 570; and Cy7, 766–855 nm for Opal 780. Regions of interest measuring 500 × 500 μm were acquired across the PFC, targeting cortical layers 1 and 2/3.

To improve spatial resolution and signal-to-noise ratio for puncta detection, raw images were deconvolved using Huygens Essential software (Scientific Volume Imaging). Image registration, cell segmentation, and transcript quantification were performed using a pipeline adapted from the previously published FijiFISH-based workflow described by Sullivan et al. (50). Cell-associated ROIs were generated using a nuclear-centric approach in which DAPI-positive nuclei served as primary seeds and were expanded by 3 μm to capture surrounding cytoplasmic mRNA puncta. The resulting ROIs were used to quantify transcript signal within each defined cell-associated region.

For each cell-associated ROI, *Egfp*, *Hmgcr*, and *Fam107a* expression were quantified as percent area covered. GFP-positive cells were defined based on the EGFP signal and classified as transfected neurons. Neighboring GFP-negative cells were defined as EGFP-negative cell-associated ROIs located within the local GFP-positive transfection hotspot. These measurements were used to compare transcript expression between GFP-positive neurons and neighboring GFP-negative cells across GFP control and C4-OE tissue.

### PLIN2 immunostaining and DAPI-associated lipid droplet quantification

#### Tissue preparation

P21–P23 mice were transcardially perfused with 1× PBS followed by 4% PFA in PBS. Brains were extracted, postfixed in 4% PFA for 24 h, transferred to 30% sucrose in PBS, and stored at 4°C until sectioning. Coronal brain sections were cut at 50 μm thickness using a Cuttec S freezing-stage sliding microtome.

#### Immunostaining and lipid droplet labeling

Free-floating coronal sections containing the medial prefrontal cortex were processed for PLIN2 immunostaining. Sections were washed three times in 1× PBS for 10 min each at room temperature on a gentle rocker. Sections were then blocked and permeabilized for 1 h at room temperature in permeabilization/blocking buffer containing 5% bovine serum albumin (BSA; Sigma-Aldrich, cat. A4737), 2% normal goat serum (NGS; Invitrogen, cat. 50062Z), and 0.5% Triton X-100 (LabChem, cat. LC262801) in 1× PBS.

Rabbit anti-PLIN2 primary antibody (Abcam, cat. AB52356; 1:500) was diluted in permeabilization/blocking buffer and applied to free-floating sections for approximately 48 h at 4°C on a rocker. After primary antibody incubation, sections were washed three times in 1× PBS for 10 min each. Sections were then incubated with Alexa Fluor 594-conjugated goat anti-rabbit IgG (H+L) secondary antibody (Life Technologies, cat. A11012; 1:200) diluted in permeabilization/blocking buffer for 2 h at room temperature, protected from light. After secondary antibody incubation, sections were washed three times in 1× PBS for 10 min each at room temperature.

Sections were mounted onto SuperFrost Plus glass slides (Thermo Fisher Scientific, cat. 26902) and allowed to dry flat, protected from light. Sections were coverslipped with SlowFade Glass Soft-Set Antifade Mountant (Invitrogen, cat. S36917) using #1.5 coverslips (Ted Pella Inc., cat. 260154), and coverslip edges were sealed with nail polish. Slides were stored at 4°C in the dark. For the present analysis, the channels used were DAPI, GFP, and PLIN2/LD signal.

#### Confocal imaging

Confocal z-stacks were acquired on a 3i Marianas inverted spinning-disk confocal microscope (Intelligent Imaging Innovations/3i) using SlideBook 6 software. Images were collected with a Zeiss 63× objective, 1.4 NA oil-immersion Plan-Apochromat, and included DAPI, GFP, and PLIN2/LD channels. The microscope was equipped with a Yokogawa CSU-X1 spinning-disk unit, an Andor iXon 888 EMCCD camera, and a piezo Z-axis stage for z-stack acquisition.

Imaging parameters were kept consistent across samples. DAPI, GFP, and PLIN2 channels were acquired using standard exposure times of 30 ms, 100 ms, and 300 ms, respectively. Confocal z-stacks were acquired with actual z-step sizes of 0.2 μm or 0.333 μm. Measurements were performed on calibrated images using a pixel size of 0.165 μm/pixel, corresponding to 6.0577 pixels/μm.

For each mouse, 2–4 imaging ROIs were selected within the GFP-positive transfection hotspot, spanning approximately 300–500 μm along the rostrocaudal axis and centered primarily within the mPFC. ROIs were selected to capture the GFP-positive transfected region and neighboring GFP-negative cells within the local transfected field, while avoiding tissue damage, imaging artifacts, or regions with poor signal quality.

#### Image analysis and PLIN2/LD puncta quantification

Image processing and quantification were performed in Fiji/ImageJ (National Institutes of Health, Bethesda, MD, USA) using custom-written macros for z-projection generation, channel-specific masking, ROI-based quantification of PLIN2/LD, and LD burden categorization. Statistical analyses and graph generation were performed in GraphPad Prism (GraphPad Software, Boston, MA, USA) and R (R Foundation for Statistical Computing, Vienna, Austria). Final figure assembly was performed in CorelDRAW (Corel Corporation, Ottawa, ON, Canada).

For each field of view, the cortical L2/3 region was manually defined to exclude pia/L1 and non-analyzable tissue regions. The same L2/3 region of interest was applied to the DAPI and PLIN2/LD channels to generate matched L2/3-only stacks. Poor-quality regions, tissue artifacts, or imaging defects were excluded prior to downstream analysis when present.

Ten-slice maximum-intensity projections were generated from the usable imaging depth for each L2/3-only stack. Because the actual z-step sizes were 0.2 μm or 0.333 μm, each 10-slice projection represented approximately 2.00–3.33 μm of optical depth. This projection strategy provided a consistent sampling unit within L2/3 while limiting excessive compression of signal across the full z-stack. Z-planes with reduced image quality, loss of intensity, blurring, or heterogeneous tissue appearance were excluded from downstream analysis when necessary. The same z-range was applied to the matched DAPI and PLIN2/LD channels for each ROI.

DAPI-positive nuclear ROIs were segmented from the DAPI channel using auto-thresholding and processed using Binary Close, Fill Holes, and Watershed to improve nuclear segmentation and separate adjacent nuclei. DAPI nuclear ROIs were identified using Analyze Particles with a size range of 30–250 μm² and circularity of 0.20–1.00.

PLIN2/LD puncta were segmented from the PLIN2 channel using the corresponding 10-slice projections. PLIN2/LD puncta were segmented using auto-thresholding, most commonly the Triangle or IsoData methods. Thresholding was selected after visual QC to identify discrete PLIN2-positive puncta while minimizing background haze, speckle overcalling, and clumping. These LD segmentation approaches were used across both experimental groups and were not preferentially assigned to one condition. PLIN2/LD puncta were identified using Analyze Particles with a size range of 0.03–30 μm² and circularity of 0.00–1.00.

Within the GFP-positive L2/3 transfection hotspot, PLIN2/LD puncta were visually enriched near DAPI-positive nuclei rather than being uniformly distributed throughout the surrounding tissue. Because cell-boundary markers were not included in this analysis, DAPI-associated ROIs were used as a reproducible spatial anchor to quantify nucleus-associated or perinuclear PLIN2/LD burden. For the main analysis, LD burden was quantified in GFP-negative DAPI-associated ROIs located within the GFP-positive transfection hotspot, representing non-transfected neighboring cells surrounding the sparsely transfected population. Accordingly, measurements are reported as DAPI-associated/perinuclear PLIN2-positive LD burden and should not be interpreted as whole-cell LD content.

For each z-block, the matching DAPI ROI set and PLIN2/LD puncta mask were used to quantify PLIN2/LD puncta within each DAPI-associated ROI. The analysis exported one row per DAPI ROI, including DAPI ROIs with zero PLIN2/LD puncta. For each DAPI ROI, the following measurements were calculated: DAPI ROI area in μm², PLIN2/LD puncta count, total PLIN2/LD puncta area in μm², PLIN2/LD puncta density normalized to DAPI ROI area, and PLIN2/LD area fraction. LD puncta density was calculated as PLIN2/LD puncta count divided by DAPI ROI area and is reported as LDs/μm². LD area fraction was calculated as total PLIN2/LD puncta area divided by DAPI ROI area and is reported as a fraction or percentage of the DAPI-associated ROI area.

For categorical analysis, each DAPI-associated ROI was assigned to an LD burden category based on PLIN2/LD puncta count: 0 LDs, 1 LD, 2–5 LDs, 6–10 LDs, 11–15 LDs, 16–20 LDs, 21–30 LDs, or >30 LDs. The percentage of DAPI-associated ROIs in each category was calculated for each imaging ROI. Additional summary metrics included the percentages of LD-negative, LD-positive, and high-LD-burden DAPI ROIs, defined as DAPI ROIs containing >10 PLIN2/LD puncta.

ROI-level results were used for visualization and quality control. Animal-level summaries were used for final group-level comparisons as described in the Statistical analysis section.

#### Statistical analysis

Statistical analyses were performed using GraphPad Prism (GraphPad Software, LLC, Boston, MA, USA) and R statistical software (R Foundation for Statistical Computing, Vienna, Austria). Data are shown as mean ± SEM unless otherwise indicated. For bulk RNA-seq, differential-expression analysis was performed using DESeq2, with significance determined by Wald tests followed by Benjamini–Hochberg FDR correction, as described above.

For MFISH analyses, Hmgcr and Fam107a expression were quantified as percent area covered within cell-associated ROIs. Cell-level measurements were summarized within each animal, and statistical analyses were performed on animal-level summary values; individual cells were not treated as independent biological replicates. Comparisons across genotype and cell category were analyzed using two-way ANOVA followed by Šídák’s multiple comparisons test.

For PLIN2/LD analyses, animal-level summaries were used as the primary biological replicates for final group-level comparisons. DAPI-associated ROIs were treated as nested within imaging ROIs and animals. LD puncta density, LD area fraction, percentage of LD-positive DAPI-associated ROIs, percentage of LD-negative DAPI-associated ROIs, and percentage of high-LD-burden DAPI-associated ROIs were compared between groups using the statistical tests indicated in the figure legends. For categorical LD burden distributions, the percentage of DAPI-associated ROIs in each LD-count category was calculated for each imaging ROI and summarized at the animal level when used for group comparisons.

For cell-type-aware DEG projection and correlation-domain analyses, the analyses were descriptive and were not used to estimate changes in cell-type abundance. DEG-defined PCA projections, PC coordinate-binning, and correlation heatmaps were used to visualize cell-type-associated transcriptional structure and gene–gene correlation domains represented within directional C4-OE bulk RNA-seq signatures. Statistical tests, significance thresholds, and exact sample definitions for quantitative analyses are reported in the corresponding figure legends.

## Results

### Experimental Design and Analysis Workflow

We employed an unbiased omics and gene-transfer approach (**Fig. 1**) to investigate whether elevated levels of the mouse homolog of the SCZ risk gene *C4A*—complement component 4b (*C4b* or *C4*)—disrupt brain function through previously unrecognized molecular mechanisms. By selectively overexpressing *C4* in a limited subset of neurons in the mPFC (9,30), we generated a sparse model of C4 dysregulation to examine how localized perturbation influences gene-expression programs across surrounding cortical cell populations (**Fig. 1A, B**).

**Figure 1.**
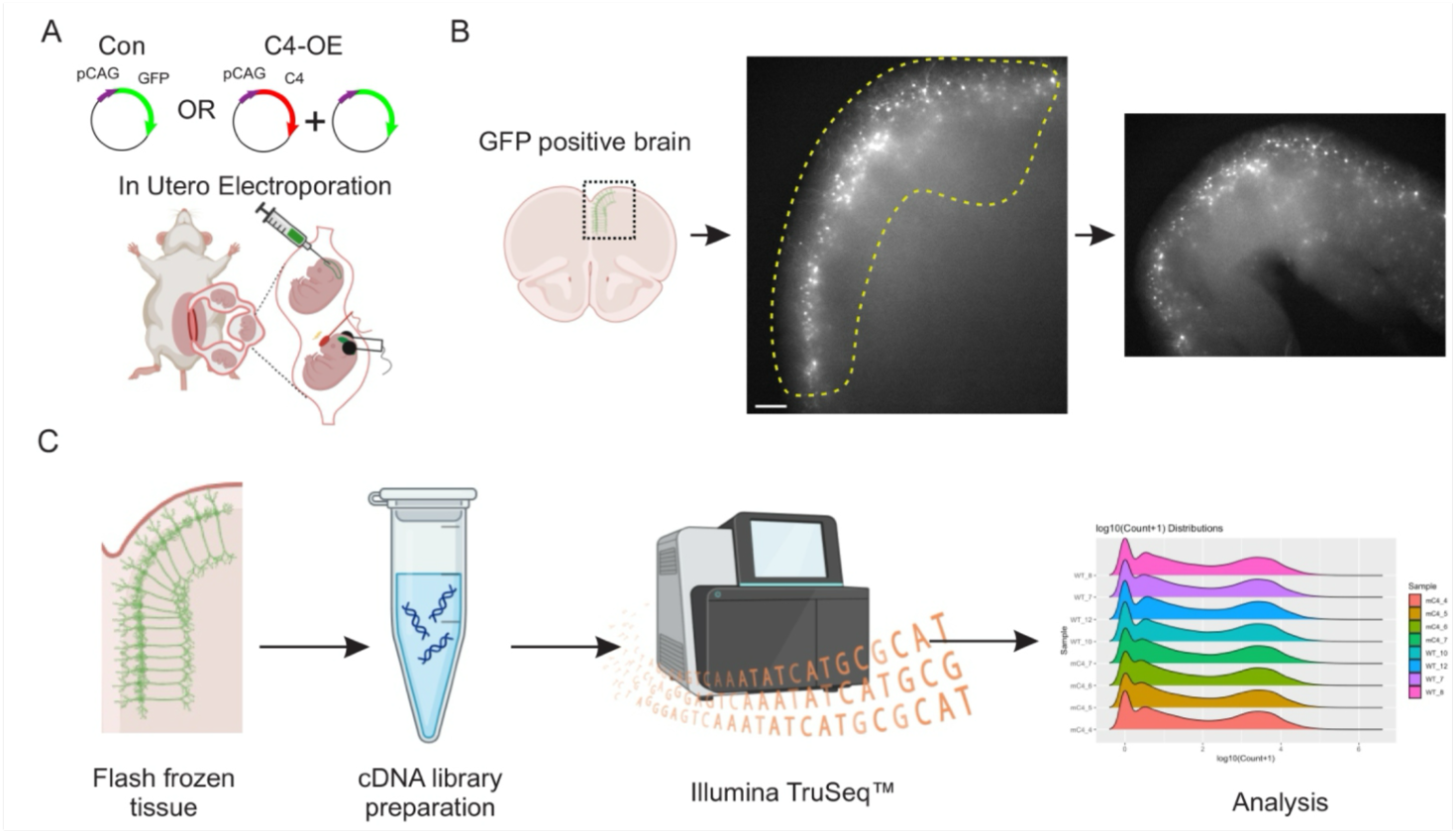
Experimental Design and Analysis Workflow. (**A**) IUE was performed on gestation day 16 (E16) CD-1 dams. Cartoon adapted from Phadke et al (https://creativecommons.org/licenses/by-nc-nd/4.0/). In control conditions, embryos received pCAG-EGFP at 1 μg/μl. In the C4-OE condition, embryos received pCAG-EGFP at 1 μg/μl together with pCAG-mC4/C4b at 1 μg/μl, corresponding to a GFP:C4 plasmid ratio of 1:1. (**B**) Cartoon and slices of the P21-23 PFC containing cells that have been isolated using microdissection (yellow dotted line). Scale bar = 100 μm. (**C**) After microdissection, mPFC tissue was immediately flash-frozen on dry ice. Next, the Broad Genomics Platform prepared cDNA libraries using the Illumina TruSeq Stranded mRNA Sample Preparation Kit (isolating at least 250 ng of purified total RNA). Whole-transcriptome sequencing was performed at the Broad Institute Genomic Services, and R (v4.2.1) was used for subsequent analysis.

Using M-FISH and super-resolution imaging, we previously demonstrated that developing L2/3 pyramidal neurons in the PFC express low baseline levels of *C4*, which localizes to synapses (9,17). Electroporation of a plasmid encoding *C4* under the control of the CAG promoter significantly increased total and synaptic C4 levels in these cortical neurons (9). Consistent with these observations, bulk RNA-seq of microdissected tissue confirmed increased *C4b* transcript abundance in C4-OE samples relative to GFP-control samples (log₂FC = 2.53, padj = 4.9 × 10⁻³). Our previous findings further showed that, aside from alterations in connectivity, C4-OE did not measurably affect the overall morphology or intrinsic excitability of transfected neurons, consistent with an absence of overt toxicity associated with increased *C4* expression (9,17). At postnatal day 21–23 (P21–23), electroporated tissue was harvested and the PFC was rapidly microdissected under fluorescence microscopy (**Fig. 1B**).

We selected the third postnatal week for microdissection for three complementary reasons. First, this developmental window is consistent with our previous studies (9,17), enabling direct comparison across datasets. Second, early postnatal development represents a critical period of synaptic maturation and refinement of cortical circuitry (30,51–53). Consistent with this, Comer et al. (9) reported a marked expansion of excitatory synapses during this period, with dendritic spine density increasing approximately 6–8-fold between P7 and P21. Third, because SCZ is a neurodevelopmental disorder (11,51,53), this early postnatal window provides an opportunity to capture molecular and circuit-level alterations associated with disease risk during a period of active cortical development.

In both control and C4-OE conditions, the isolated PFC tissue contained transfected pyramidal neurons together with surrounding non-transfected neuronal and non-neuronal cell populations. Based on the anatomical distribution of GFP fluorescence (**Fig. 1B**), the microdissected cortical slab encompassed layers L1, L2/3, L5, and L6 of the medial PFC. To estimate IUE transfection efficiency, we quantified GFP-positive neurons in the PFC at P21 across six sections comprising 108 ROIs and an estimated total volume of ∼2.43 mm³ (9,54). Approximately 4,000 GFP-positive neurons were identified (9,54), corresponding to ∼1.4–2.4% of the estimated 164,000– 286,000 neurons within this volume (33). Thus, IUE produced the expected sparse neuronal labeling, with the overwhelming majority of cells within the analyzed tissue remaining non-transfected.

To assess the broader molecular consequences of sparse neuronal C4 elevation, we next analyzed transcriptome-wide gene-expression changes in microdissected mPFC tissue (**Fig. 1C**).

### Sparse Neuronal C4 Overexpression is Associated with Broad Transcriptional Remodeling in the mPFC

To investigate the transcriptional impact of sparse neuronal C4 overexpression in the mPFC, we performed differential expression analysis comparing C4-OE and control samples using DESeq2 (35), with sex included as an additive covariate in the model (∼ sex + genotype; Methods). RNA-seq libraries showed adequate sequencing depth across samples, and we observed the expected relationship between gene-level mean expression and variance (**S1 Fig. A, B**). Sample sex assignments were independently confirmed by the reciprocal expression patterns of *Xist* and the Y-linked gene *Ddx3y* (**S1 Fig. C**). Principal component analysis of variance-stabilized expression values showed separation of C4-OE and control samples primarily along PC1, which explained 33% of the variance, whereas PC2 explained 18% and captured additional sex-associated variation among control samples (**S1 Fig. D**).

Differential expression analysis identified 1,573 genes with a Benjamini–Hochberg-adjusted P value < 0.05, including 849 upregulated and 724 downregulated genes in C4-OE tissue (**S1 Table**). These changes reflected a broad, bidirectional transcriptional response (**Fig. 2A**). Hierarchical clustering of the 50 genes with the smallest adjusted P values further illustrated distinct expression patterns across C4-OE and control samples (**Fig. 2B**).

**Figure 2.**
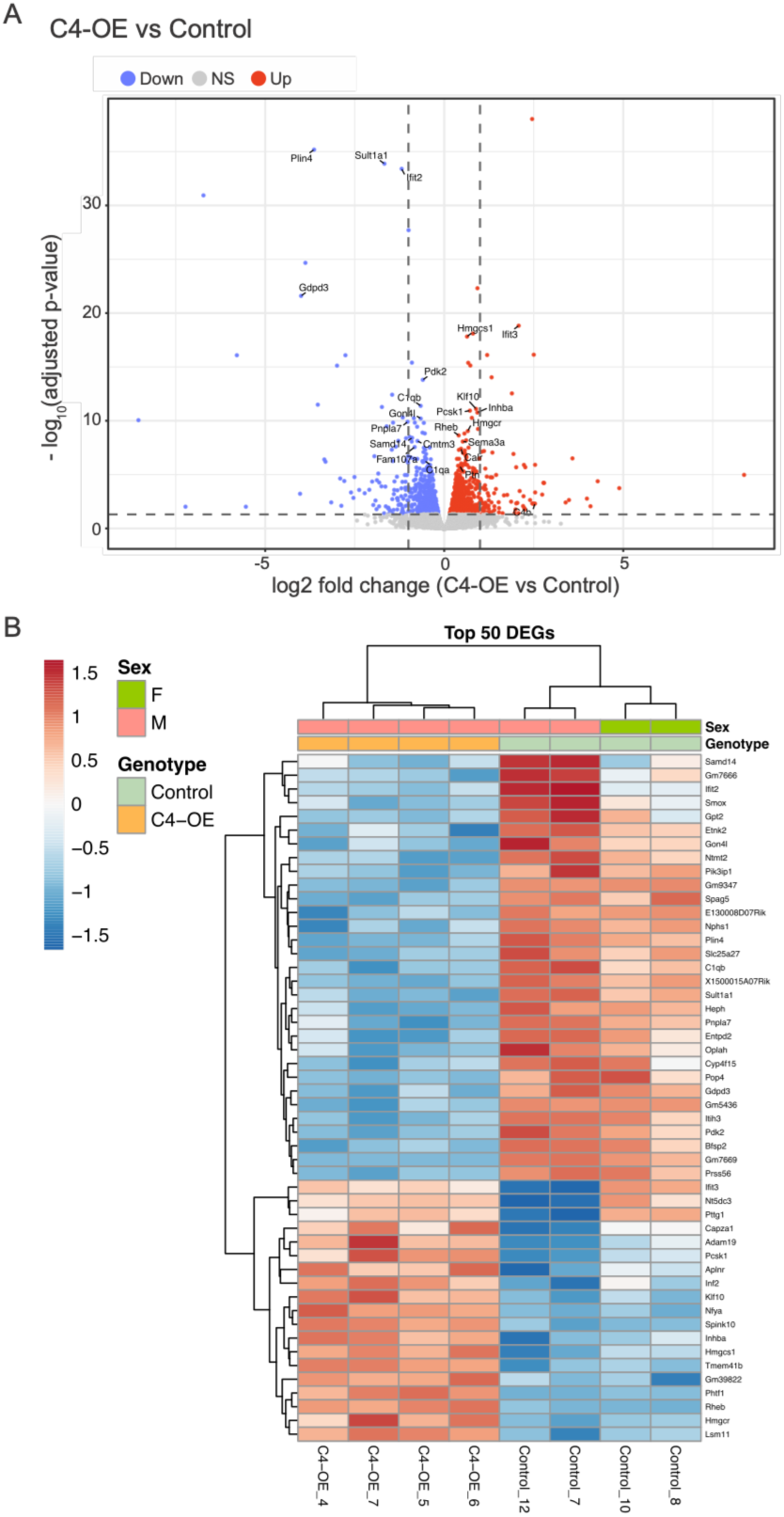
Sparse neuronal C4 elevation induces gene-level transcriptional changes associated with lipid metabolism, complement signaling, and synaptic plasticity. **(A**) Volcano plot of differential gene expression between C4-OE and Control cortical samples. Each point represents a gene; the x-axis shows DESeq2 log₂ fold change for C4-OE versus Control and the y-axis shows −log₁₀ adjusted P value. Genes with adjusted P < 0.05 are shown in red when upregulated and blue when downregulated, whereas nonsignificant genes are shown in gray. Dashed vertical lines at log₂ fold change ±1 are included as visual references and were not used to define differential expression. Selected genes associated with lipid metabolism, complement/immune signaling, synaptic plasticity, and cellular stress are labeled. (**B**) Heatmap of the 50 genes with the smallest DESeq2 adjusted P values. DESeq2-normalized counts were log₂-transformed after addition of a pseudocount of 1 and Z-scored independently for each gene across samples. Columns represent individual samples and are annotated by genotype and sex; rows and columns were hierarchically clustered. Colors indicate gene-wise Z-scored expression using a display range of −1.5 to +1.5 Z.

*C4b*, the experimentally overexpressed transcript, was significantly increased in C4-OE tissue (log₂FC = 2.53; adjusted P = 4.93 × 10⁻³; baseMean = 3,265; **S1 Fig. E**), consistent with successful detection of the intended perturbation despite the sparse transfection strategy. The differential expression results were not driven predominantly by low-abundance transcripts: none of the 1,573 significant DEGs had a DESeq2 baseMean below 20, and only 13 genes (0.83%) had either a baseMean below 20 or a mean normalized expression below 20 in control samples.

### Gene-level Transcriptional Changes Identify Lipid, Complement, and Plasticity-associated Responses

A prominent feature of the transcriptional response was the coordinated upregulation of genes involved in cholesterol biosynthesis, including *Cyp51*, *Mvd*, *Hmgcr*, *Sqle*, *Nsdhl*, *Msmo1*, *Sc5d*, *Dhcr7*, *Idi1*, and *Hmgcs1* (**S1 Table**). Although the effect sizes of individual genes were generally modest, their consistent direction across multiple steps of the pathway suggested coordinated regulation of cholesterol biosynthesis-associated transcription. Notably, *Hmgcs1* and *Hmgcr* were among the 50 genes with the smallest adjusted P values and displayed consistent expression differences between C4-OE and control samples (**Fig. 2B**). *Hmgcr*, which encodes the rate-limiting enzyme in cholesterol biosynthesis and has been implicated as a modifier of neurological disease risk (55–57), was significantly upregulated (**S1 Table**).

Several of the most significant DEGs were also associated with lipid-droplet biology, phospholipid metabolism, and lipid turnover (58). *Plin4*, the second most significant DEG, was strongly downregulated and was among the prominent genes identified in the differential-expression analysis (**Fig. 2A, B**). *Gdpd3*, *Pnpla7*, *Etnk2*, and *Cyp4f15* were also significantly reduced, while *Plin3* was significantly downregulated but was not among the 50 genes with the smallest adjusted P values (**S1 Table**). Together with the coordinated increase in cholesterol biosynthesis-associated transcripts, these findings identified lipid metabolism and lipid-droplet-associated gene expression as prominent components of the C4-OE transcriptional response. These observations motivated subsequent examination of lipid-droplet burden within the C4-OE cortical field.

Complement- and immune-associated transcripts exhibited changes in both directions. In contrast to the experimentally driven increase in *C4b*, all three genes encoding the C1q complex— *C1qa, C1qb,* and *C1qc*—were significantly downregulated (**S1 Table**). Additional immune-associated genes, including *Ccl4*, *Tlr1*, *Hc*, and *Chia1*, were also reduced, whereas the MHC class II-associated transcript *H2-Ea* was strongly upregulated (**S1 Table**). Thus, sparse neuronal C4 elevation was associated with a heterogeneous complement- and immune-related transcriptional response rather than a uniform directional shift across inflammatory genes. Because these measurements were obtained from bulk cortical tissue, the cellular sources and functional consequences of these expression changes could not be resolved directly.

Genes associated with synaptic plasticity, cellular growth, and homeostatic stress responses were also significantly altered. These included upregulation of *Homer1* (59,60), an immediate-early gene and postsynaptic scaffolding protein involved in regulating synaptic strength and excitability; *Rheb* (61,62), a positive regulator of mTOR signaling and cellular growth; and *Klf10* (63), an activity-responsive transcription factor (**Fig. 2A, B; S1 Table**). *Calr* was also upregulated, whereas *Itih3* and *Heph* were downregulated (**S1 Table**). These changes identified additional transcriptional responses involving genes associated with synaptic plasticity, growth signaling, immune regulation, cellular stress, and metal homeostasis.

At the gene level, the C4-OE transcriptional response showed several prominent features, including coordinated upregulation of cholesterol biosynthesis-associated genes, altered lipid-metabolism and lipid-droplet-associated transcripts, downregulation of all three C1q genes, and changes in genes linked to synaptic plasticity and cellular stress. We therefore next asked whether these individual transcriptional changes converged on broader biological pathways and processes and subsequently examined their spatial and cellular relevance within the surrounding cortical microenvironment.

### Convergent Transcriptional Programs Following Sparse C4-OE

To identify coordinated biological programs associated with sparse C4 elevation, we applied two complementary enrichment approaches to the DESeq2 output: gene set enrichment analysis (GSEA) on the full ranked gene list and Gene Ontology (GO) over-representation analysis (ORA) on the significant DEGs using all tested genes as the background universe. Because ORA evaluates a thresholded subset of the ranked list analyzed by GSEA in full, agreement between the two approaches indicates that a biological signal is robust to the choice of significance cutoff rather than representing independent replication.

### Cholesterol and Sterol Biosynthesis Programs are Positively Enriched Following Sparse C4-OE

One of the strongest and most consistent transcriptional signals involved cholesterol and sterol biosynthesis. GSEA identified positive enrichment of cholesterol-related gene sets across multiple collections, including WP CHOLESTEROL BIOSYNTHESIS (NES = +2.49, adjusted P = 6.2 × 10⁻⁷), REACTOME CHOLESTEROL BIOSYNTHESIS (NES = +2.36), HALLMARK CHOLESTEROL HOMEOSTASIS (NES = +1.97), and GOBP STEROL BIOSYNTHETIC PROCESS (NES = +1.95) (**Fig. 3A, B; S2–S4 Tables**). GO ORA of the upregulated DEGs identified related biological processes, including cholesterol biosynthetic process via desmosterol and via lathosterol (11 genes each; adjusted P = 1.0 × 10⁻⁵), zymosterol biosynthetic and metabolic processes (8 genes; adjusted P = 7.4 × 10⁻⁴), and steroid biosynthetic process (20 genes) (**Fig. 3C; S5 Table**).

**Figure 3.**
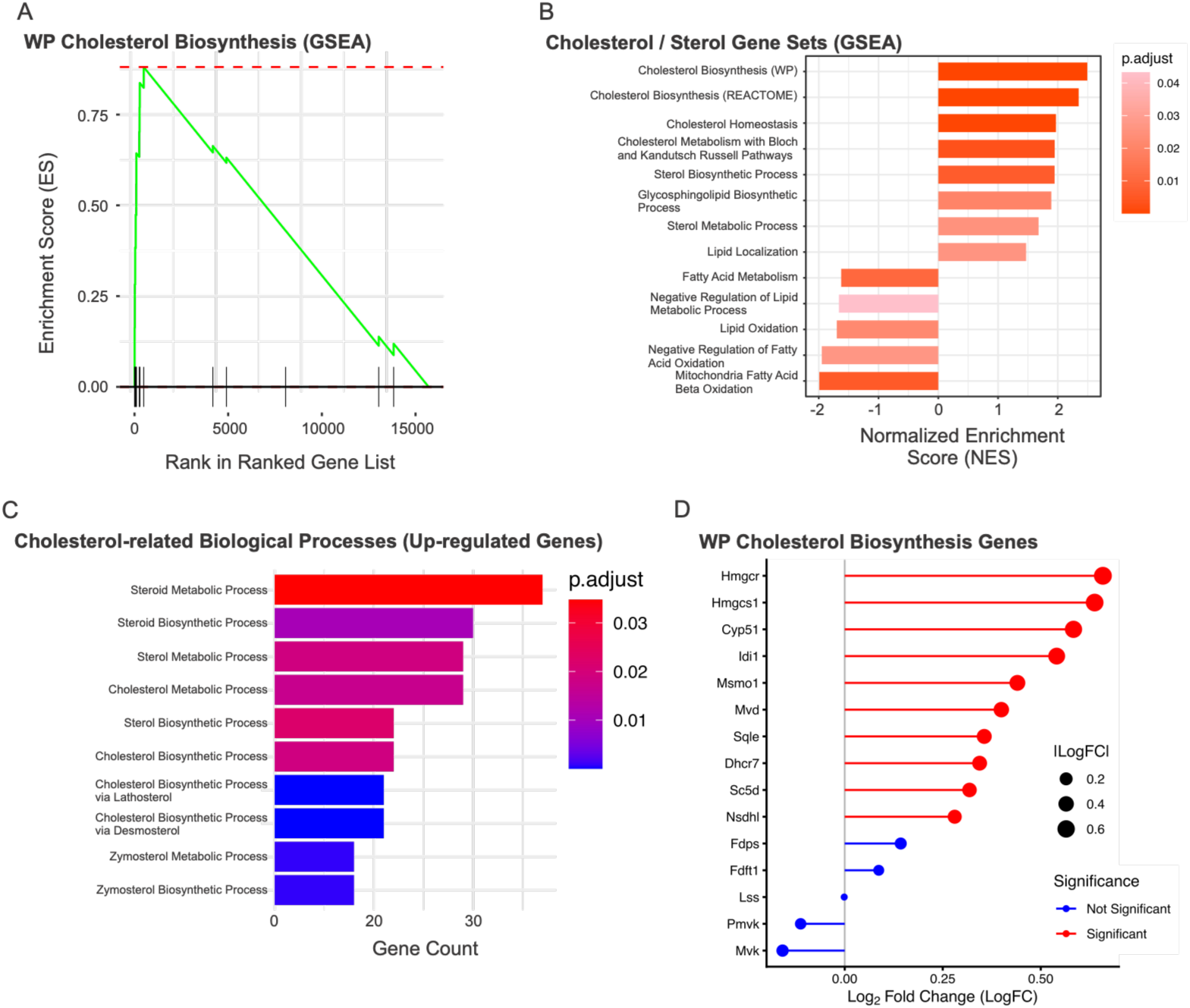
Sparse neuronal C4 elevation is associated with coordinated enrichment of cholesterol and sterol biosynthesis programs. (**A**) Gene set enrichment analysis (GSEA) enrichment plot for the WikiPathways cholesterol biosynthesis gene set, showing the running enrichment score across the ranked DESeq2 gene list. (**B**) Selected cholesterol-, sterol-, and lipid-associated gene sets identified by GSEA. Bars indicate normalized enrichment score (NES), with positive values representing enrichment toward genes increased in C4-OE and negative values representing enrichment toward genes decreased in C4-OE. Bar color indicates FDR-adjusted P value. Only gene sets meeting FDR-adjusted P < 0.05 are shown. (**C**) Gene Ontology Biological Process over-representation analysis of upregulated DEGs showing cholesterol-, sterol-, and steroid-related biological processes. Bar length indicates the number of genes contributing to each term, and color indicates adjusted P value. Only terms meeting adjusted P < 0.05 are shown. (**D**) Gene-level DESeq2 log₂ fold changes for genes in the WikiPathways cholesterol biosynthesis gene set. Red points indicate genes significantly differentially expressed at adjusted P < 0.05 and blue points indicate nonsignificant genes; point size represents absolute log₂ fold change.

At the gene level, 10 of the 15 genes in the WikiPathways cholesterol biosynthesis set were significantly differentially expressed, and all 10 were upregulated: *Hmgcs1*, *Hmgcr*, *Cyp51*, *Idi1*, *Msmo1*, *Mvd*, *Sqle*, *Dhcr7*, *Sc5d*, and *Nsdhl* (**Fig. 3D; S1 Table**). Individual effect sizes were modest (log₂FC 0.28–0.66), but the changes were distributed across multiple steps of the pathway. These included *Hmgcs1* (log₂FC = +0.64, adjusted P = 1.5 × 10⁻¹⁸) and *Hmgcr* (log₂FC = +0.66, adjusted P = 9.0 × 10⁻¹⁰), together with several downstream sterol-modifying enzymes. Consistent with broader changes in lipid-associated transcription, GO molecular function analysis of upregulated genes also identified lipoprotein particle binding (*Lrp8*, *Sorl1*, *Lpl*, *Ldlr*, *Pltp*, *Lsr*, and *Scarb1*; adjusted P = 3.4 × 10⁻²; **S6 Table**).

GSEA additionally identified negative enrichment of fatty acid metabolism (NES = −1.63, adjusted P = 1.0 × 10⁻²) and mitochondrial fatty acid β-oxidation (NES = −1.97, adjusted P = 7.8 × 10⁻³; **Fig. 3B; S3 Table**). Overall, these results indicate coordinated transcriptional regulation of multiple lipid-associated pathways, with positive enrichment of cholesterol biosynthesis programs accompanied by negative enrichment of fatty acid oxidation-related programs.

### Synaptic Organization Programs are Positively Enriched while Synaptic Translation Programs are Negatively Enriched

A second prominent transcriptional pattern involved synaptic and axonal programs. GO ORA of upregulated DEGs identified regulation of synapse organization (47 genes; adjusted P = 1.3 × 10⁻³), synapse assembly (34 genes), axonogenesis (49 genes), axon extension (19 genes), and inhibitory synapse assembly (7 genes) (**Fig. 4A; S5 Table**). Cellular component analysis further identified enrichment of genes annotated to the postsynaptic membrane (44 genes; adjusted P = 6.5 × 10⁻³), postsynaptic specialization membrane (25 genes), dendritic shaft (12 genes), and proximal dendrite (5 genes), together with additional membrane and extracellular compartments (**Fig. 4B; S7 Table**).

**Figure 4.**
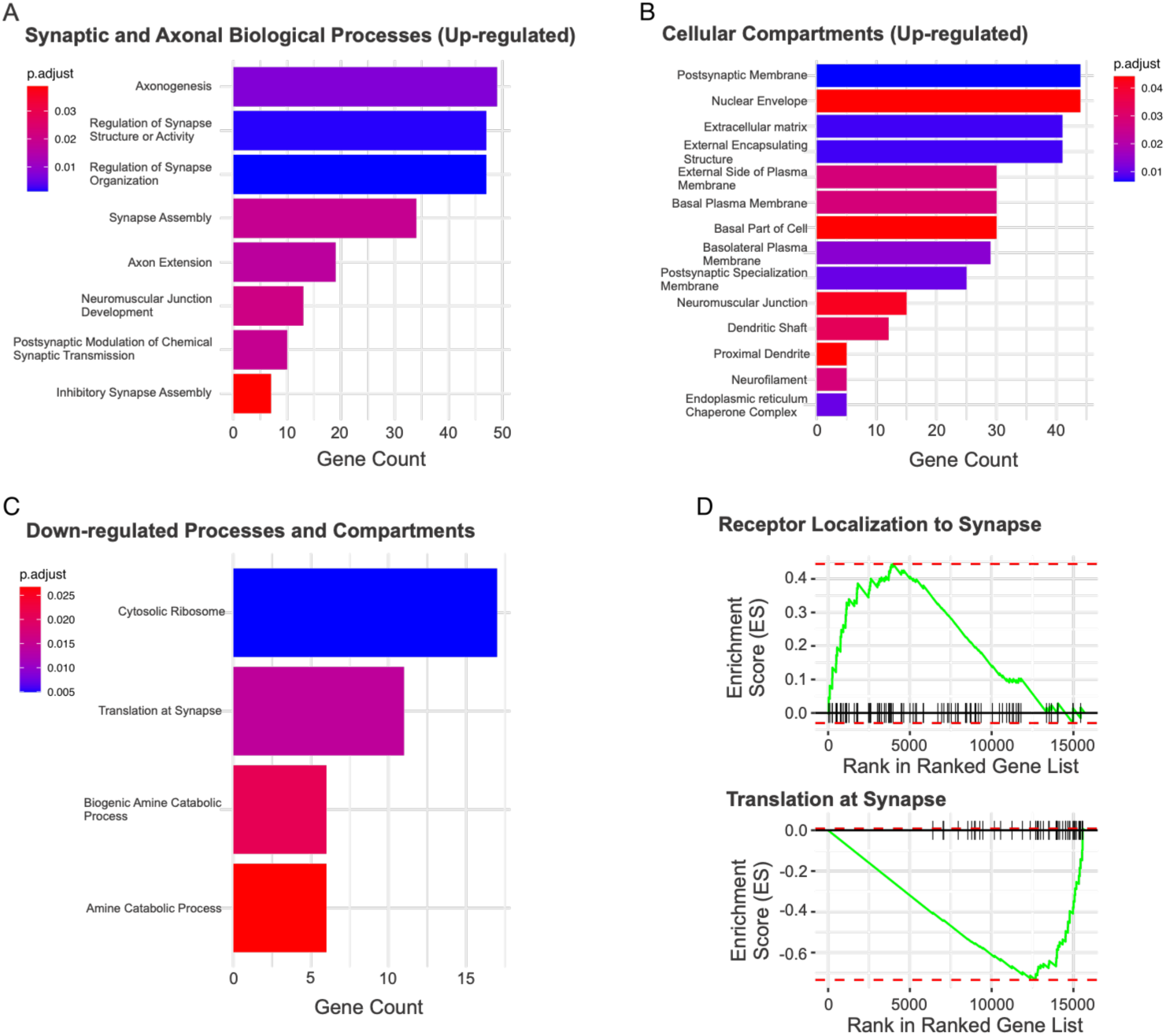
Sparse neuronal C4 elevation is associated with opposing synaptic organization and translation-related transcriptional programs. (**A**) Selected Gene Ontology (GO) Biological Process terms enriched among upregulated DEGs, highlighting synaptic organization, synapse assembly, and axonal development-related processes. Bar length indicates the number of genes contributing to each term, and color indicates adjusted P value. Only terms meeting adjusted P < 0.05 are shown. (**B**) Selected GO Cellular Component terms enriched among upregulated DEGs, including postsynaptic and dendritic compartments. Bar length indicates gene count, and color indicates adjusted P value. Only terms meeting adjusted P < 0.05 are shown. (**C**) Selected GO Biological Process and Cellular Component terms enriched among downregulated DEGs, including translation at synapse and the cytosolic ribosome. Bar length indicates gene count, and color indicates adjusted P value. Only terms meeting adjusted P < 0.05 are shown. (**D**) Representative GSEA enrichment plots illustrating positive enrichment of receptor localization to synapse (NES = +1.83) and negative enrichment of translation at synapse (NES = −2.77, FDR-adjusted P = 7.3 × 10⁻⁹) across the ranked DESeq2 gene list. Vertical ticks indicate the positions of genes belonging to each gene set within the ranked list.

GSEA supported the same direction of change, with positive enrichment for axon development (NES = +1.82, adjusted P = 1.0 × 10⁻⁶), postsynapse organization (NES = +1.52), receptor localization to synapse (NES = +1.83), regulation of presynapse organization (NES = +1.81), and positive regulation of dendrite development (NES = +1.90; **Fig. 4D; S4 Table**).

In contrast, the downregulated DEG set returned relatively few significant terms, but these were concentrated in translational and ribosomal processes. GO ORA identified translation at synapse (11 genes; adjusted P = 1.4 × 10⁻²) and cytosolic ribosome (17 genes; adjusted P = 5.0 × 10⁻³) (**Fig. 4C; S8** and **S9 Tables**). GSEA showed stronger negative enrichment of related programs: GOBP TRANSLATION AT SYNAPSE was among the most negatively enriched GO biological process gene sets (NES = −2.77, adjusted P = 7.3 × 10⁻⁹), while canonical pathway analysis identified cytoplasmic ribosomal proteins (NES = −2.91, adjusted P = 2.3 × 10⁻¹⁴), nonsense-mediated decay (NES = −2.67), and eukaryotic translation initiation (NES = −2.58) among the most negatively enriched pathways (**Fig. 4D; S3** and **S4 Tables**). The leading-edge genes in the translation-at-synapse set consisted predominantly of ribosomal protein genes together with *Eef2*. Thus, sparse C4 elevation was associated with opposing transcriptional patterns involving positive enrichment of synaptic organization-related programs and negative enrichment of synaptic translation-related programs.

### Vascular, Adhesion, and Inflammatory Programs are Enriched Following Sparse C4-OE

A third transcriptional theme involved vascular, endothelial, adhesion, and extracellular-matrix-associated programs. GO ORA of upregulated genes identified endothelial cell migration among the most significantly enriched processes (36 genes; adjusted P = 2.0 × 10⁻⁵), together with blood vessel endothelial cell migration (23 genes), regulation of vasculature development (34 genes), regulation of angiogenesis (33 genes), sprouting angiogenesis (19 genes), endothelium development (20 genes), and establishment of endothelial barrier (11 genes) (**Fig. 5A; S5 Table**). Consistent with these findings, GSEA identified HALLMARK ANGIOGENESIS as a strongly positively enriched Hallmark gene set (NES = +2.24, adjusted P = 3.0 × 10⁻⁵; **Fig. 5D; S2 Table**).

**Figure 5.**
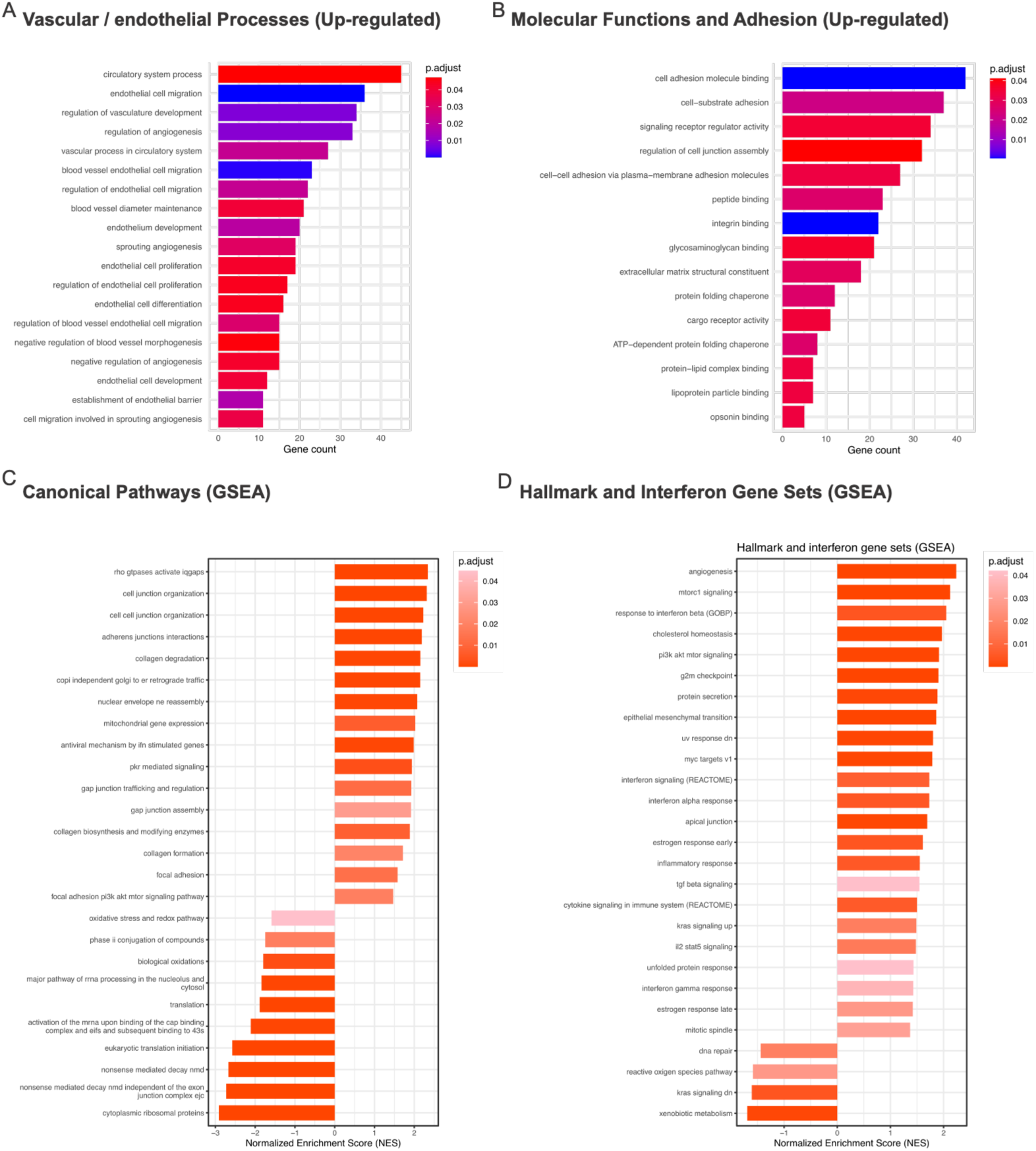
Sparse neuronal C4 elevation is associated with enrichment of vascular, adhesion, extracellular-matrix, and inflammatory transcriptional programs. (**A**) Selected Gene Ontology (GO) Biological Process terms enriched among upregulated DEGs, highlighting vascular and endothelial-related programs, including endothelial cell migration, regulation of vasculature development, angiogenesis, and endothelial barrier-related processes. Bar length indicates the number of genes contributing to each term, and color indicates adjusted P value. Only terms meeting adjusted P < 0.05 are shown. (**B**) Selected GO Biological Process and Molecular Function terms enriched among upregulated DEGs, highlighting adhesion- and extracellular-matrix-associated programs, including cell–substrate adhesion, cell–cell adhesion via plasma-membrane adhesion molecules, regulation of cell junction assembly, cell adhesion molecule binding, integrin binding, and extracellular matrix structural constituent. Bar length indicates gene count, and color indicates adjusted P value. Only terms meeting adjusted P < 0.05 are shown. (**C**) Selected canonical pathway gene sets identified by GSEA. Positively enriched pathways include adhesion-, junction-, and extracellular-matrix-related programs, whereas negatively enriched pathways include translation- and RNA-processing-related programs. Bars indicate normalized enrichment score (NES), with positive values representing enrichment toward genes increased in C4-OE and negative values representing enrichment toward genes decreased in C4-OE. Bar color indicates FDR-adjusted P value. Only gene sets meeting FDR-adjusted P < 0.05 are shown. (**D**) Selected Hallmark and interferon-related gene sets identified by GSEA, highlighting angiogenesis, interferon/inflammatory-response, signaling, and additional transcriptional programs. Bars indicate normalized enrichment score (NES), with positive values representing enrichment toward genes increased in C4-OE and negative values representing enrichment toward genes decreased in C4-OE. Bar color indicates FDR-adjusted P value. Only gene sets meeting FDR-adjusted P < 0.05 are shown.

Adhesion- and extracellular-matrix-associated terms were also enriched. GO molecular function analysis of upregulated genes identified cell adhesion molecule binding (42 genes; adjusted P = 4.8 × 10⁻⁴), integrin binding (22 genes; adjusted P = 4.8 × 10⁻⁴), and extracellular matrix structural constituent (18 genes), while biological process analysis identified cell–substrate adhesion (37 genes), cell–cell adhesion via plasma-membrane adhesion molecules (27 genes), and regulation of cell junction assembly (32 genes) (**Fig. 5B; S5** and **S6 Tables**). Canonical pathway GSEA similarly identified positive enrichment of cell junction organization (NES = +2.31, adjusted P = 4.6 × 10⁻⁶), cell–cell junction organization (NES = +2.23), adherens junction interactions (NES = +2.19), and collagen degradation (NES = +2.15) (**Fig. 5C; S3 Table**). Because sparse C4-OE was restricted to a small fraction of excitatory neurons, the emergence of these vascular, endothelial, adhesion, and matrix-associated transcriptional programs is consistent with transcriptional responses extending beyond the directly transfected neuronal population.

GSEA also identified positive enrichment of interferon- and inflammatory-response gene sets, including response to interferon beta (NES = +2.05, adjusted P = 4.2 × 10⁻³), REACTOME INTERFERON SIGNALING (NES = +1.73), HALLMARK INTERFERON ALPHA RESPONSE (NES = +1.73), HALLMARK INFLAMMATORY RESPONSE (NES = +1.55), and cytokine signaling in the immune system (NES = +1.50) (**Fig. 5D; S2** and **S3 Tables**). Because sex and genotype were partially confounded in the experimental design, we compared leading-edge genes with genes independently identified as sex-differential. Sex-associated genes were restricted to 3 of 16 leading-edge genes in the response-to-interferon-beta set (*Ifit1, Ifit3,* and *Iigp1*) and were absent from the broader interferon, cytokine, and inflammatory leading-edge sets, indicating that these broader enrichment signals were not driven primarily by the genes flagged as sex-differential.

Additional positively enriched Hallmark programs included mTORC1 signaling (NES = +2.12), PI3K–AKT–mTOR signaling (NES = +1.91), G2M checkpoint (NES = +1.90), epithelial–mesenchymal transition (NES = +1.86), and MYC targets (NES = +1.79; **Fig. 5D; S2 Table**). Given the predominantly post-mitotic state of the P21 cortex, we interpret these pathway labels as transcriptional signatures rather than evidence of neuronal proliferation. Negative enrichment was also observed for several Hallmark pathways, including DNA repair (NES = −1.44), reactive oxygen species pathway (NES = −1.59), and xenobiotic metabolism (NES = −1.69; **Fig. 5D; S2 Table**). Molecular function analysis of downregulated genes did not identify significantly enriched terms (**S10 Table**).

Together, these analyses reveal a broad transcriptional response to remarkably sparse neuronal C4 elevation. Despite direct manipulation of only ∼1.4–2.4% of neurons within the sampled cortical volume, C4-OE was associated with coordinated changes in cholesterol and sterol biosynthesis, synaptic organization and translation-related programs, and vascular, adhesion, extracellular-matrix, and inflammatory signatures across the surrounding mPFC tissue. Thus, sparse neuronal C4 elevation is associated with widespread cortical transcriptional remodeling, consistent with a response that extends beyond the directly manipulated neuronal population.

### WGCNA Identifies a Prominent Cholesterol Biosynthesis-associated Co-expression Program

To determine whether the transcriptional changes associated with sparse neuronal C4 elevation were organized into coordinated gene networks, we performed weighted gene co-expression network analysis (WGCNA) (44). Using a soft-thresholding power of 15, the network achieved a signed scale-free topology fit of R² = 0.833 and resolved 25 non-grey co-expression modules (**S11 Table**). We Z-scored module eigengenes across samples to visualize module-level expression patterns (**Fig. 6A**).

**Figure 6.**
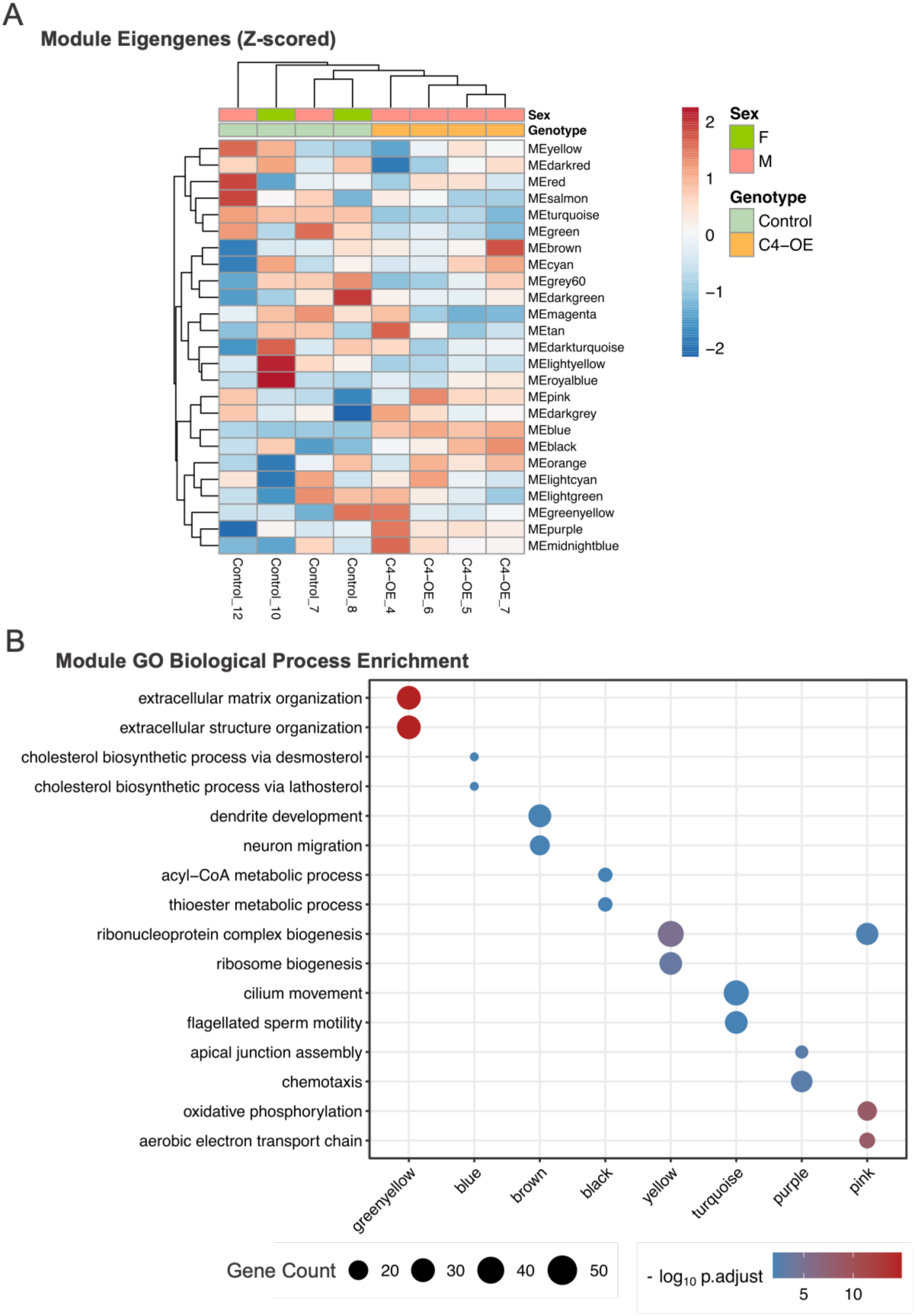
WGCNA identifies a cholesterol biosynthesis-associated co-expression program following sparse neuronal C4 elevation. (**A**) Heatmap of Z-scored module eigengenes identified by weighted gene co-expression network analysis (WGCNA) using a soft-thresholding power of 15. Columns represent individual samples and are annotated by genotype and sex; rows represent the 25 non-grey co-expression modules. Module eigengenes and samples are hierarchically clustered to visualize module-level expression patterns across the dataset. The blue module showed the strongest positive association with genotype, whereas the turquoise module showed a significant association in the opposite direction. (**B**) Selected Gene Ontology (GO) Biological Process terms significantly enriched among genes assigned to representative WGCNA modules. The x-axis indicates module identity and the y-axis indicates enriched biological processes. Point size represents the number of module genes contributing to each GO term, and point color represents −log₁₀ adjusted P value.

The blue module showed the strongest association with genotype (r = 0.993, P = 9.8 × 10⁻⁷, adjusted P = 2.5 × 10⁻⁵; **S17 Table**), with all four Control samples and all four C4-OE samples exhibiting opposing module-level expression patterns (**Fig. 6A**). The turquoise module was also significantly associated with genotype, but in the opposite direction (r = −0.985, adjusted P = 1.1 × 10⁻⁴; **S17 Table**), and was enriched primarily for cilium- and motility-associated biological processes (**Fig. 6B; S18 Table**). No other module remained significantly associated with genotype after multiple-testing correction. We interpreted these module–genotype associations in the context of the partial confounding between sex and genotype described above.

GO biological process enrichment revealed that the blue module was strongly enriched for cholesterol biosynthesis. Its two most significantly enriched processes were cholesterol biosynthetic process via desmosterol and cholesterol biosynthetic process via lathosterol, each represented by 11 genes (adjusted P = 9.8 × 10⁻³; **Fig. 6B; S18 Table**). These genes included *Acat2, Cyp51, Dhcr7, Ebp, Hmgcr, Hmgcs1, Msmo1, Mvd, Nsdhl, Sc5d,* and *Sqle*. Notably, nine of the ten cholesterol-biosynthesis genes highlighted in the preceding differential-expression analysis were assigned to the blue module (**S15** and **S16 Tables**).

This network-level organization reinforced the cholesterol signature identified in the preceding gene-level and pathway-enrichment analyses. Sparse neuronal C4 was associated with coordinated upregulation of multiple cholesterol-biosynthesis genes, while both GSEA and GO over-representation analysis identified positive enrichment of cholesterol and sterol biosynthetic programs, including the same desmosterol- and lathosterol-associated processes (**Fig. 3; S1–S5 Tables**). Thus, C4 elevation was associated not only with changes in individual cholesterol-associated transcripts, but also with a coordinated cholesterol-associated co-expression pattern spanning multiple steps of the biosynthetic pathway.

Other modules were enriched for distinct biological programs (**Fig. 6B; S17** and **S18 Tables**), including extracellular matrix and extracellular structure organization, dendrite development and neuron migration, acyl-CoA and thioester metabolism, ribosome and ribonucleoprotein biogenesis, apical junction assembly and chemotaxis, and oxidative phosphorylation and aerobic electron transport. These processes overlapped with several of the broader transcriptional responses identified in the preceding pathway analyses, indicating that sparse neuronal C4 elevation was associated with coordinated changes across multiple biological programs.

We next evaluated the sensitivity of the WGCNA results to the selected soft-thresholding power. Across powers 12–18, signed scale-free topology fit ranged from R² = 0.780 to 0.860, and the networks consistently resolved 24–26 non-grey modules (**S11–S15 Tables**). Individual gene-to-module assignments varied substantially across network solutions (median adjusted Rand index = 0.31), indicating that gene-level module membership should be interpreted with caution. The blue module at the reference power of 15 was among the more stable modules across alternative network solutions, with the second-highest module stability score (**S14 Table**). This sensitivity analysis supported the selection of power 15 as a representative network solution rather than identifying a uniquely optimal power.

Together, these findings identify cholesterol biosynthesis as a prominent component of the C4-OE-associated transcriptional response. Despite direct C4 elevation in only a small fraction of cortical neurons, cholesterol-related signals were detected across differential-expression, pathway-enrichment, and co-expression network analyses. Although gene-level WGCNA module assignments varied across alternative network solutions, the convergence of these complementary analyses supports cholesterol biosynthesis as one of the more consistent transcriptional features associated with sparse neuronal C4 elevation.

### Sparse Neuronal C4 Elevation Shows Selective Molecular Correspondence with the Schizophrenia Synaptic Proteome

Our previous studies, including Comer et al. and Phadke et al. (9,17), demonstrated that increased neuronal C4 levels produce specific synaptic alterations, including reduced excitatory connectivity and impaired AMPA receptor trafficking. Given the genetic association of *C4* with SCZ, we therefore asked whether the molecular consequences of sparse neuronal C4 elevation engage pathways that are also altered at synapses in SCZ. To address this, we compared the mouse C4-OE cortical transcriptome with a synaptic proteomic dataset from individuals with SCZ. After restricting the analysis to high-confidence one-to-one human–mouse orthologs measurable in both datasets, 7,498 genes constituted the shared cross-species universe (**Fig. 7A**).

**Figure 7.**
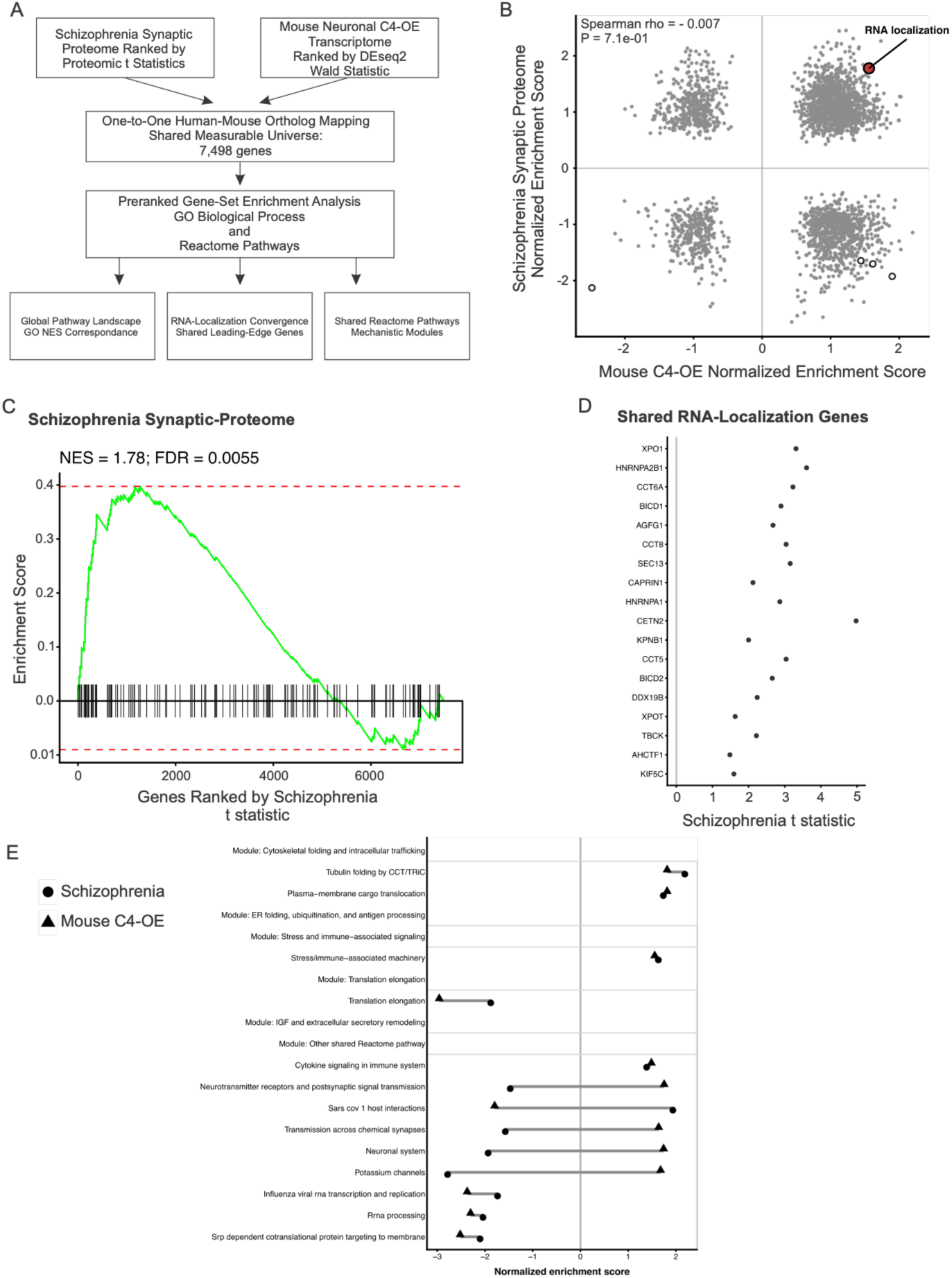
Sparse neuronal C4 elevation shows selective molecular correspondence with the schizophrenia synaptic proteome. (**A**) Cross-species analysis workflow comparing the SCZ synaptic proteome with the mouse neuronal C4-OE transcriptome. Proteins in the SCZ dataset were ranked by proteomic t statistic and mouse genes by the DESeq2 Wald statistic. High-confidence one-to-one human–mouse ortholog mapping defined a shared measurable universe of 7,498 genes used for preranked gene set enrichment analysis of Gene Ontology (GO) Biological Process and Reactome pathways. Analyses assessed global pathway correspondence, RNA-localization leading-edge overlap, and shared Reactome pathways. (**B**) Comparison of normalized enrichment scores (NES) for 2,739 GO Biological Process gene sets tested in both datasets. The x-axis shows mouse C4-OE NES and the y-axis shows SCZ synaptic-proteome NES. Global pathway correspondence was minimal (Spearman ρ = −0.007, P = 0.71). Hollow circles indicate the four pathways significant at FDR < 0.05 in both datasets; RNA localization is highlighted in red. (**C**) GSEA enrichment plot for the GO Biological Process RNA-localization gene set in the SCZ synaptic proteome (NES = +1.78, FDR = 0.0055). RNA localization also showed positive enrichment in the mouse C4-OE transcriptome (NES = +1.56, nominal P = 0.0034), although this signal did not remain significant after multiple-testing correction (FDR = 0.126). Vertical ticks indicate the positions of RNA-localization genes within the ranked SCZ proteomic dataset. (**D**) Shared RNA-localization leading-edge genes identified independently in the schizophrenia synaptic proteome and mouse C4-OE transcriptome. The two leading-edge sets contained 40 and 46 genes, respectively, with 18 genes shared between datasets. The plot shows the SCZ proteomic t statistic for these shared genes. (**E**) Reactome pathways significantly enriched at FDR < 0.05 in both datasets. Circles indicate SCZ synaptic-proteome NES values and triangles indicate mouse C4-OE transcriptome NES values; connecting lines link the values for each pathway. Thirteen pathways were significant in both datasets and included both same-direction and opposite-direction enrichment across programs related to translation and RNA processing, protein folding and intracellular trafficking, immune/stress signaling, and neuronal and synaptic function.

We first assessed whether the two datasets showed broad pathway-level correspondence. Across GO Biological Process terms, enrichment scores were essentially uncorrelated between the mouse C4-OE transcriptome and the SCZ synaptic proteome (Spearman ρ = −0.007, P = 0.71; **Fig. 7B**), indicating little global similarity in pathway-level responses. Of 2,739 GO biological processes tested in both datasets, 190 reached FDR < 0.05 in the SCZ synaptic proteome and 46 in the mouse C4-OE transcriptome, but only four were significant in both. This overlap was not greater than expected by chance (4 observed versus 3.19 expected; Fisher P = 0.397). Moreover, three of the four shared pathways showed opposite enrichment directions. Thus, the overall GO analysis supported limited global correspondence between the C4-OE transcriptome and the SCZ synaptic proteome.

Against this background, RNA localization showed a notable same-direction signal. RNA localization was positively enriched in the SCZ synaptic proteome (NES = +1.78, FDR = 0.0055) and showed positive enrichment in the mouse C4-OE transcriptome (NES = +1.56; nominal P = 0.0034), although the mouse enrichment did not remain significant after multiple-testing correction (FDR = 0.126; **Fig. 7B, C**). Thus, RNA localization was significantly enriched in the SCZ synaptic proteome and showed a similar positive enrichment trend in the C4-OE transcriptome, despite little overall correspondence between the two pathway landscapes.

Leading-edge analysis identified additional overlap within this pathway. The SCZ and mouse RNA-localization enrichments contained 40 and 46 leading-edge genes, respectively, with 18 genes shared between the two sets (**Fig. 7D**; Shared RNA-Localization Genes). The shared genes were *AGFG1, AHCTF1, BICD1, BICD2, CAPRIN1, CCT5, CCT6A, CCT8, CETN2, DDX19B, HNRNPA1, HNRNPA2B1, KIF5C, KPNB1, SEC13, TBCK, XPO1,* and *XPOT*. These genes include components associated with RNA and nucleocytoplasmic transport, intracellular trafficking, and protein-folding machinery, providing a shared leading-edge core underlying the positive RNA-localization enrichment in the two datasets. The analysis defined this core as the intersection of the independently derived leading edges.

We next asked whether selective correspondence was also evident using Reactome pathways. Thirteen Reactome pathways reached FDR < 0.05 in both datasets (**Fig. 7E**). Several showed the same enrichment direction, including rRNA processing, SRP-dependent cotranslational protein targeting to membrane, eukaryotic translation elongation, formation of tubulin-folding intermediates by CCT/TRiC, GLUT4 translocation, adaptive immune system, and cytokine signaling. In contrast, potassium channels, neuronal system, transmission across chemical synapses, and neurotransmitter receptor/postsynaptic signaling pathways showed opposite enrichment directions. Shared Reactome pathways were grouped for biological interpretation into broader categories including cytoskeletal folding and intracellular trafficking, ER folding and antigen processing, translation elongation, IGF and extracellular secretory remodeling, and stress- and immune-associated signaling. These findings again indicated that correspondence was concentrated within selected biological programs rather than broadly distributed across the two datasets.

Gene-level comparisons showed a similarly limited pattern. Using SCZ-associated synaptic proteins at FDR < 0.10 and mouse C4-OE DEGs at adjusted P < 0.05, 43 orthologous genes overlapped compared with 36.2 expected by chance, which was not significant by Fisher exact test (P = 0.128). When the SCZ threshold was restricted to FDR < 0.05, 23 genes overlapped compared with 12.4 expected, corresponding to a 1.86-fold enrichment (P = 0.0023). Thus, gene-level overlap was detectable under the more stringent SCZ threshold, but the magnitude was modest and dependent on the significance criterion. Directional concordance among overlapping genes was not broadly increased, further arguing against a uniform molecular shift shared between C4-OE and SCZ.

Overall, these analyses indicate that sparse neuronal C4 elevation does not broadly reproduce the molecular landscape of the SCZ synaptic proteome. Instead, the datasets show limited global correspondence with selective overlap in specific molecular programs. RNA localization provides one such example, showing positive enrichment in both datasets and an 18-gene shared leading-edge core, while concordant Reactome pathways identify additional points of correspondence involving intracellular trafficking, RNA and protein processing, translation, and immune-associated signaling.

### Directional C4-OE Signatures Are Associated with Distinct Cortical Reference Programs

The preceding analyses showed that sparse neuronal C4-OE is associated with broad transcriptional remodeling in the local cortical field, including altered expression of genes associated with cholesterol biosynthesis, neuronal development, axon guidance, synaptic plasticity, vascular remodeling, and *C4b*-associated co-expression networks (**Figs. 2–6**). However, because bulk RNA-seq was performed from microdissected cortical tissue containing transfected neurons embedded within surrounding non-transfected cells, these transcriptional changes could reflect multiple cortical cell-state-associated programs. We therefore next asked whether the directional C4-OE-regulated DEG signatures mapped onto defined cortical transcriptional landscapes.

To do this, we leveraged a reference-based analytical framework (46) using the Allen Brain Map Mouse Whole Cortex and Hippocampus SMART-seq single-cell dataset and its associated 10x– SMART-seq taxonomy (64). We curated this reference to retain cortical/isocortical populations and collapsed transcriptomic subclasses into major neuronal and non-neuronal cortical cell classes for projection of C4-OE-regulated bulk RNA-seq signatures (**S2 Fig. A–D**). After excluding hippocampal populations, the curated reference retained cells from 16 cortical/isocortical regions and 31 transcriptomic subclasses, which were consolidated into 19 grouped cortical cell classes for downstream analysis (**S19–S21 Tables**). Dimensionality reduction of the curated reference using PCA, t-SNE, and UMAP resolved major cortical neuronal, glial, immune, and vascular populations, while marker-gene analysis confirmed expected broad cell-type identities (**S2 Fig. A–D**). These analyses established a reference framework for contextualizing C4-OE-regulated bulk RNA-seq signatures within cortical cell-type-associated transcriptional space.

### Directional C4-OE Signatures Map onto Distinct Astrocyte- and Vascular-Associated Reference Spaces

We then projected the 250 C4-OE-downregulated and 250 C4-OE-upregulated DEGs with the largest directional log2 fold changes, after intersection with the curated cortical reference, independently onto the reference dataset. This approach identified cell-type-associated transcriptional structure within the bulk RNA-seq signatures and was not intended to infer changes in cell-type abundance. The genes used for each directional projection are provided in **S22 and S23 Tables**.

The C4-OE-downregulated DEG signature produced a prominent separation of astrocyte-associated reference cells along PC1, indicating that genes expressed at lower levels in C4-OE tissue map prominently onto an astrocyte-associated reference-cell axis (**Fig. 8A**). Other cortical reference populations remained more closely distributed near the origin or partially overlapped along PC1. In contrast, PC2 contained additional neuronal-associated structure, with intratelencephalic excitatory populations, including L2/3 IT cells, occupying distinct regions of the projection space. Thus, the downregulated signature contained both a prominent astrocyte-associated PC1 axis and additional neuronal-associated structure along PC2 (**Fig. 8A**).

**Figure 8.**
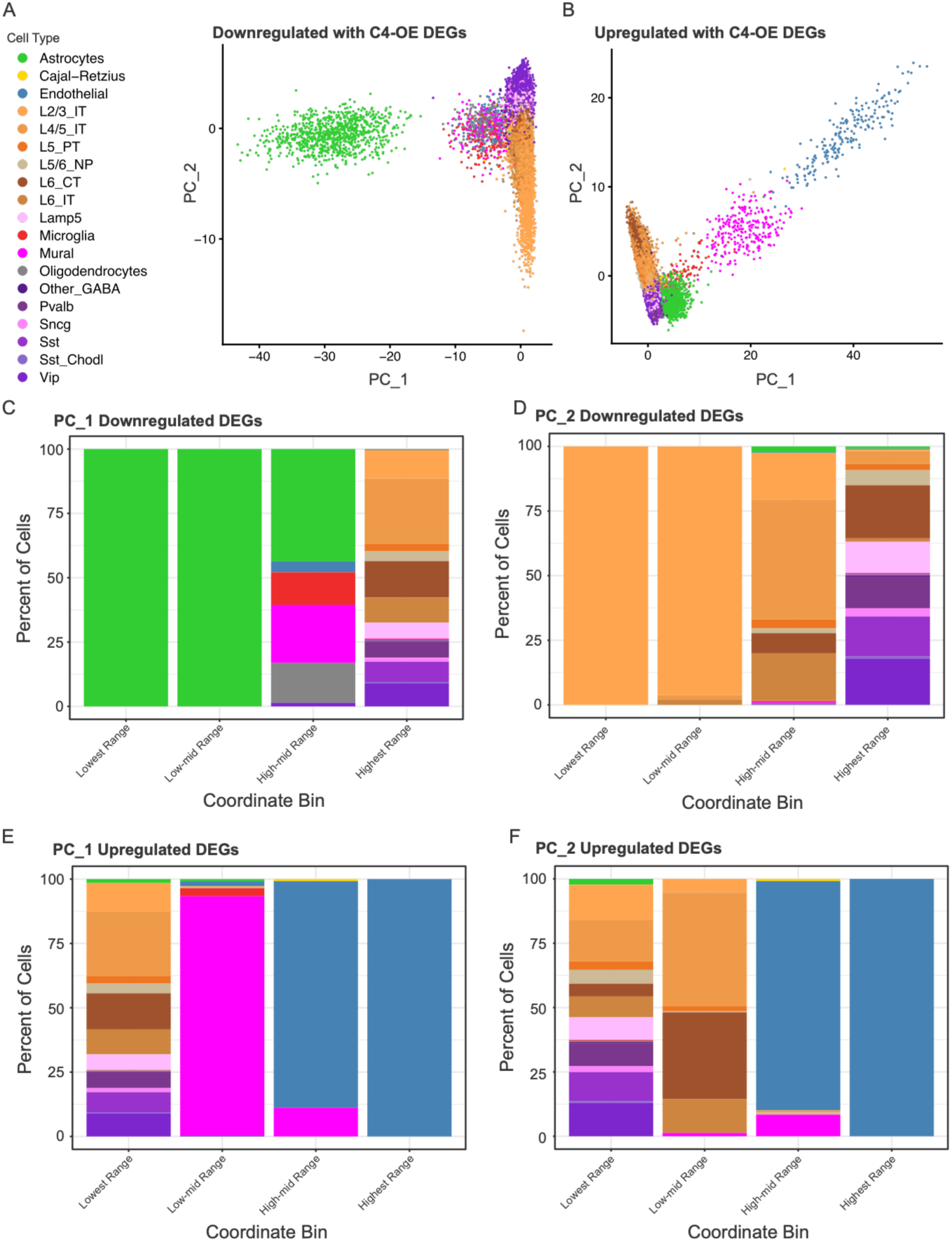
Directional C4-OE signatures map onto distinct cortical reference spaces. (**A, B**) PCA-based projection of the 250 C4-OE-downregulated (**A**) and 250 C4-OE-upregulated (**B**) DEGs with the largest directional log2 fold changes after intersection with the curated cortical scRNA-seq reference derived from the Allen Brain Map Mouse Whole Cortex and Hippocampus SMART-seq dataset. Reference cells are colored by consolidated cortical cell class. (**C–F**) PC coordinate-binning analysis showing the percentage of each cortical reference cell class across four equal-width coordinate bins for PC1 and PC2 in the downregulated (**C, D**) and upregulated (**E, F**) projections. These analyses summarize cell-type-associated structure within DEG-defined PCA space and do not estimate cell-type abundance in bulk RNA-seq samples. The genes used for the downregulated and upregulated projections are provided in **S22 and S23 Tables**, respectively. Abbreviations: C4-OE, complement C4 overexpression; DEG, differentially expressed gene; IT, intratelencephalic; L2/3, cortical layers 2/3; PC, principal component; PCA, principal component analysis; scRNA-seq, single-cell RNA sequencing.

The C4-OE-upregulated DEG signature resolved a distinct non-neuronal axis associated with vascular reference-cell structure along PC1 (**Fig. 8B**). Mural cells occupied an intermediate position along this axis, whereas endothelial cells occupied its extreme, while excitatory neuronal, interneuron, and remaining non-neuronal classes clustered more closely near the origin. PC2 also retained neuronal-associated structure, although endothelial-associated reference cells remained prominent at the upper extreme. This vascular-associated organization was consistent with the pathway-level findings from **Figures 3–6**, where C4-OE-upregulated genes were associated with angiogenesis, vascular remodeling, and structural reorganization.

Together, the downregulated and upregulated projections suggest directionally asymmetric transcriptional organization: the downregulated signature maps prominently onto an astrocyte-associated PC1 axis, whereas the upregulated signature maps onto a vascular-associated PC1 axis with mural- and endothelial-associated structure. At the same time, PC2 captures additional neuronal-associated organization, including L2/3 IT-associated structure that is particularly relevant to the IUE-targeted population.

To summarize the cell-type-associated structure across the DEG-defined principal-component axes, we divided the observed PC1 and PC2 coordinate ranges into four equal-width bins, corresponding to the lowest, low-mid, high-mid, and highest coordinate intervals, and calculated the percentage of each cortical reference cell class within each bin. In the C4-OE-downregulated projection, astrocytes accounted for the entirety of the two lowest PC1 coordinate bins, with the high-mid bin containing a mixture of astrocytes, mural cells, oligodendrocytes, microglia, and endothelial cells, and neuronal classes confined to the highest bin (**Fig. 8C**). This distribution was consistent with the prominent astrocyte-associated separation observed in the PCA projection.

PC2 binning of the downregulated projection revealed strong representation of intratelencephalic excitatory neuron classes, with L2/3 IT populations accounting for essentially all cells in the two lowest PC2 coordinate bins and deeper-layer IT and interneuron classes distributed across the higher bins (**Fig. 8D**). This L2/3-associated structure is particularly relevant to the experimental design because IUE primarily targets developing upper-layer excitatory neurons, making L2/3 IT cells the reference population most closely corresponding to the targeted neuronal population. The prominence of L2/3 IT cells along PC2 is therefore consistent with neuronal-associated transcriptional structure being most prominently represented in the reference class most closely related to the manipulated cells, although this should not be interpreted as evidence for altered L2/3 neuron abundance.

In the C4-OE-upregulated projection, PC1 binning was instead dominated by vascular-associated reference populations: mural cells comprised the large majority of the low-mid bin, endothelial cells the large majority of the high-mid bin, and endothelial cells the entirety of the highest bin, with neuronal classes restricted primarily to the lowest bin (**Fig. 8E**). PC2 binning again separated endothelial cells at the upper extreme, whereas excitatory neuronal classes occupied the lower coordinate bins (**Fig. 8F**). These binning analyses therefore reinforced the distinct organization of the two directional signatures, with astrocyte-associated structure predominating along PC1 in the downregulated projection, vascular-associated structure predominating along PC1 in the upregulated projection, and additional neuronal-associated structure captured along PC2.

Therefore, sparse neuronal C4 elevation in the mPFC is associated with directionally distinct cortical transcriptional signatures, including astrocyte-associated and vascular-associated structure along PC1 and L2/3 IT-associated neuronal structure along PC2. These findings support the idea that a sparse neuronal perturbation is associated with transcriptional signatures extending beyond the directly targeted neuronal population.

### Directional C4-OE Signatures Reveal Astrocyte-Associated Lipid/Stress and Vascular/Endothelial–Interferon, ECM, and Cholesterol Correlation Domains

We next examined gene–gene correlation structure within each directional DEG set using the cortical reference.

Among C4-OE-downregulated genes, the heatmap revealed several visually apparent correlation domains (**Fig. 9A; S24 Table**). For clarity, we annotated two prominent and biologically interpretable domains in the main figure: an astrocyte-associated lipid–metabolic stress domain (module 2) and a complement/immune-stress-response domain (module 4). The astrocyte-associated lipid–metabolic stress domain included lipid-droplet and lipid-turnover genes, such as *Plin3, Plin4,* and *Pnpla2*, as well as the SCZ-relevant metabolic gene *Prodh* (65). Notably, these genes were embedded within a larger domain containing astrocyte-associated and metabolic-support genes, including *Aldoc, Fam107a, Gjb6, Hepacam, Ntsr2,* and *Etnppl*, providing gene-level support for the astrocyte-associated structure observed in the PCA projection. The complement/immune-stress-response domain included the complement genes *C1qa, C1qb,* and *C1qc*, immune/stress-associated genes *Rgs1, Ccl4,* and *Socs3*, and the immediate-early stress-response genes *Junb* and *Gadd45g*. Remaining downregulated domain assignments are provided in **S24 Table**.

**Figure 9.**
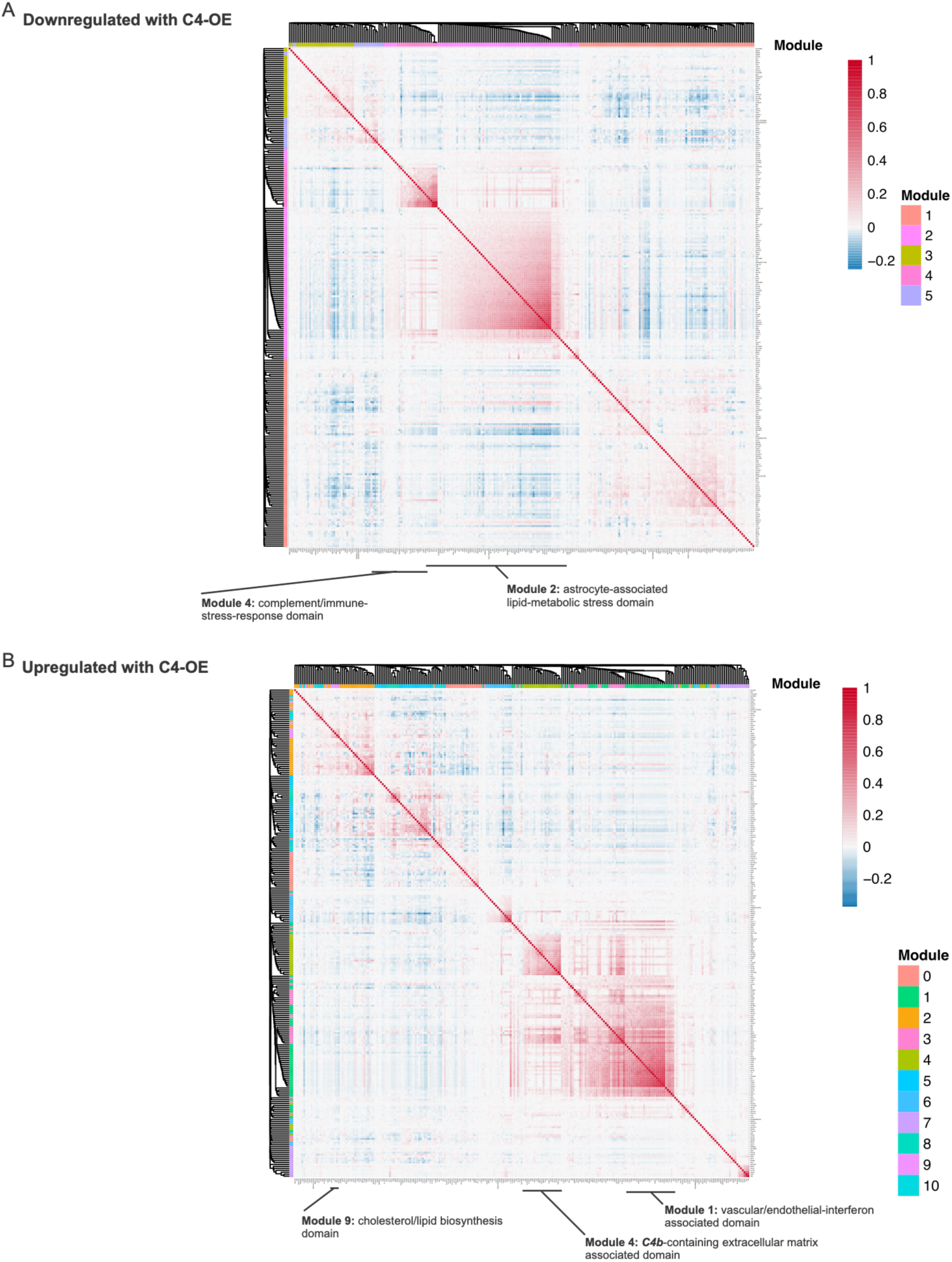
Directional C4-OE signatures reveal distinct gene–gene correlation domains in cortical reference space. Gene–gene Pearson correlation heatmaps were generated from the same directional DEG signatures used for the cortical reference projection in Fig. 8: C4-OE-downregulated (**A**) and C4-OE-upregulated (**B**). Genes with zero variance across reference cells were excluded before correlation analysis; the downregulated matrix retained all 250 genes, whereas the upregulated matrix contained 249 genes after exclusion of one zero-variance gene. Correlation matrices were hierarchically clustered, and selected correlation domains are annotated in each panel. Complete domain assignments for the downregulated and upregulated signatures are provided in **S24 and S25 Tables**, respectively. Abbreviations: C4-OE, complement C4 overexpression; DEG, differentially expressed gene.

The C4-OE-upregulated DEG heatmap also showed visually structured correlation domains (**Fig. 9B; S25 Table**). For clarity, we annotated three prominent domains in the main figure: a vascular/endothelial–interferon-associated domain (module 1), a *C4b*-containing extracellular matrix-associated domain (module 4), and a cholesterol/lipid biosynthesis domain (module 9). The vascular/endothelial–interferon-associated domain was the largest upregulated domain and contained endothelial-associated genes *Emcn, Kdr, Nos3, Cdh5,* and *Tie1* together with interferon-response genes *Isg15, Ifi44, Ifit1, Ifit3, Nos2,* and *Cd74*, consistent with the vascular-associated separation observed in the upregulated PCA projection. The *C4b*-containing extracellular matrix-associated domain contained *C4b* together with *Lbp, Lum, Fbln5,* and *Nid2*. The cholesterol/lipid biosynthesis domain included *Hmgcr, Hmgcs1, Cyp51, Idi1,* and *Stard4*, consistent with the cholesterol biosynthesis signature identified in **Figure 3**. Remaining upregulated domain assignments are provided in **S25 Table**.

Overall, these correlation-domain analyses indicate that the directional C4-OE signatures are associated with distinct gene–gene correlation domains within cortical reference space. The downregulated signature is associated with astrocyte-associated lipid/metabolic and complement/immune-stress domains, whereas the upregulated signature is associated with vascular/endothelial–interferon, *C4b*-containing extracellular matrix, and cholesterol/lipid biosynthesis domains. These findings further support the idea that sparse neuronal C4 elevation is associated with transcriptional signatures mapping onto multiple cortical cell-state-associated programs.

### Sparse C4-OE Is Associated with Non-cell-autonomous Transcriptomic Shifts in Neighboring Cortical Cells

Bulk RNA-seq differential gene expression analysis revealed that sparse C4-OE in the mPFC was associated with broad transcriptomic changes, including significant alterations in genes linked to cholesterol biosynthesis and structural/metabolic pathways. Among these C4-OE–altered genes were *Hmgcr*, which encodes a key enzyme in cholesterol biosynthesis (66), and *Fam107a*, a gene associated with cytoskeletal and metabolic regulation. Because we detected these changes in bulk tissue despite a low transfection rate, we hypothesized that focal C4 elevation may be associated with a non-cell-autonomous transcriptomic shift in the surrounding unmanipulated network. To investigate this with single-cell spatial resolution, we performed MFISH targeting *Hmgcr* and *Fam107a* in GFP control and C4-OE L2/3 tissue and quantified single-cell transcript-associated fluorescence using a previously established MFISH image-analysis pipeline (**Fig. 10A, B** (50)).

**Figure 10.**
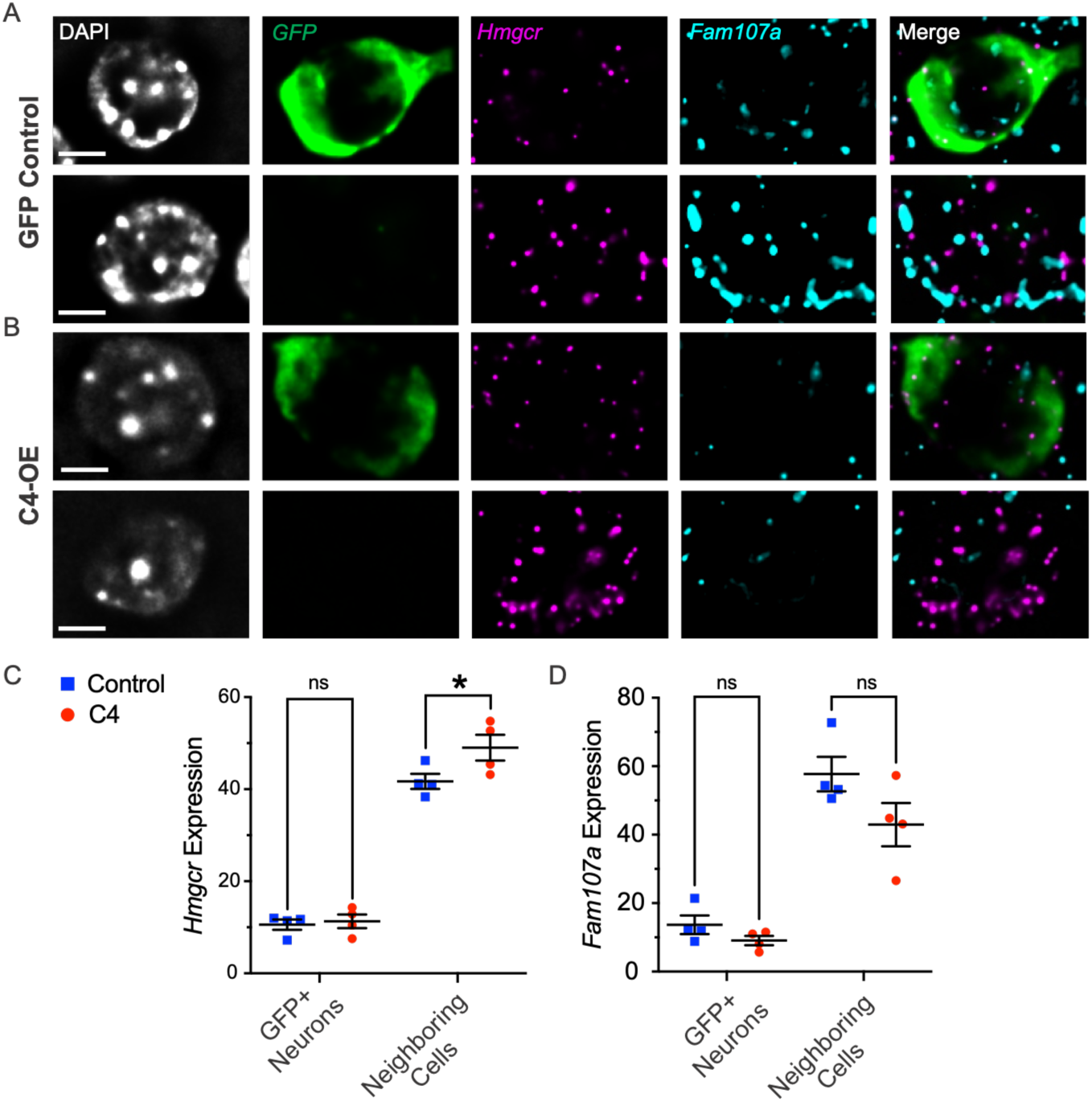
MFISH analysis of Hmgcr and Fam107a expression in GFP-positive and neighboring GFP-negative cortical cells. (**A, B**) Representative multiplex fluorescence in situ hybridization images from mPFC coronal sections of GFP control (**A**) and C4-OE (**B**) tissue. Each row shows a representative single cell imaged at 60× magnification. Channels are shown as follows: DAPI, grayscale; GFP/Opal 520, green; *Hmgcr*/Opal 570, magenta; *Fam107a*/Opal 780, cyan; and merged overlay without DAPI. Top rows show GFP-positive transfected neurons; bottom rows show neighboring GFP-negative cells. Scale bars, 5 μm. (**C**) Quantification of Hmgcr expression, measured as percent area covered, in GFP-positive neurons and neighboring GFP-negative cells from GFP control and C4-OE tissue. GFP control is shown in blue and C4-OE in red. GFP-positive neurons: GFP control, 10.59 ± 1.13%; C4-OE, 11.29 ± 1.48%. Neighboring GFP-negative cells: GFP control, 41.70 ± 1.64%; C4-OE, 49.02 ± 2.80%. Data are shown as mean ± SEM; individual points represent biological replicates; n = 4/group. Statistics: two-way ANOVA with Šídák’s multiple comparisons test; GFP-positive neurons, ns; neighboring GFP-negative cells, \**P* = 0.034. (**D**) Quantification of *Fam107a* expression, measured as percent area covered, in GFP-positive neurons and neighboring GFP-negative cells from GFP control and C4-OE tissue. GFP control is shown in blue and C4-OE in red. GFP-positive neurons: GFP control, 13.68 ± 2.70%; C4-OE, 9.08 ± 1.37%. Neighboring GFP-negative cells: GFP control, 57.71 ± 5.06%; C4-OE, 42.94 ± 6.31%. Data are shown as mean ± SEM; individual points represent biological replicates; n = 4/group. Statistics: two-way ANOVA with Šídák’s multiple comparisons test; GFP-positive neurons, ns; neighboring GFP-negative cells, *P* = 0.0637. For panels **C** and **D**, cell-level measurements were summarized within each animal, and statistical analyses were performed on animal-level summary values; individual cells were not treated as independent biological replicates.

MFISH confirmed that electroporation remained sparse, with GFP-positive cells representing approximately 7% of the total sampled cell population within the local transfection hotspot analyzed here (**S3 Fig. A**). This value is expected to be higher than our previous estimate of approximately 2% electroporation efficiency across larger mPFC regions, because those broader analyses included heterogeneous tissue areas spanning the GFP-positive hotspot, peri-transfected regions, and more sparsely labeled portions of the mPFC. To first determine whether the electroporation/transfection procedure itself influenced gene expression independently of C4, we pooled GFP-positive cells from both conditions and compared them with GFP-negative neighboring cells. GFP-positive transfected cells showed lower *Hmgcr* and *Fam107a* signal compared with GFP-negative neighboring cells, suggesting that the transfected neuronal population occupies a distinct baseline molecular state relative to the surrounding non-transfected cells (**S3 Fig. B, C**).

Building on this baseline, we next asked whether C4-OE specifically altered transcript expression in either the manipulated GFP-positive neurons or the surrounding GFP-negative neighboring cells. GFP-positive neurons showed no significant differences in *Hmgcr* or *Fam107a* signal between C4-OE and GFP control tissue (**Fig. 10C, D**). In contrast, neighboring GFP-negative cells in C4-OE tissue exhibited altered transcript expression compared with equivalent neighboring cells in GFP control tissue. Specifically, neighboring GFP-negative cells showed a significant increase in *Hmgcr* signal and a strong trend toward decreased *Fam107a* signal (**Fig. 10C, D**). These findings indicate that, for the genes examined here, the most prominent C4-OE-associated expression changes were detected in neighboring GFP-negative cells rather than in the directly transfected neurons, supporting a non-cell-autonomous component to the local transcriptional response.

These MFISH data support a model in which sparse neuronal C4-OE is associated with spatially localized transcriptomic remodeling in neighboring GFP-negative cells. This provides single-cell spatial support for the bulk RNA-seq findings and suggests that focal neuronal C4 elevation can influence molecular states within the surrounding cortical network.

#### C4-OE Increases Perinuclear Lipid-droplet Burden in Neighboring L2/3 Cells

Bulk RNA-seq revealed significant dysregulation of genes associated with lipid-droplet (LD) biology and lipid turnover, including *Plin3, Plin4, Pnpla2 (ATGL)*, and *Pnpla7*. Because LDs are increasingly implicated in CNS disease and perilipin proteins regulate LD stability and turnover (67,68), we next assessed LD burden using PLIN2 immunostaining, a widely used marker of LD accumulation in aging and neurodegenerative disease (69,70).

Representative confocal maximum-intensity projections from L2/3 showed PLIN2-positive LD puncta throughout the local transfection field in both GFP control and C4-OE tissue (**Fig. 11A, B**). In control tissue, PLIN2/LD signal was present at relatively low levels around DAPI-labeled nuclei, with scattered puncta in GFP-positive and neighboring GFP-negative regions (**Fig. 11A**). In contrast, C4-OE tissue showed more prominent PLIN2/LD signal associated with perinuclear regions within the GFP-positive transfection hotspot and in adjacent GFP-negative neighboring cells (**Fig. 11B**). Orthogonal XY, YZ, and XZ views further supported the close spatial association of PLIN2-positive puncta with DAPI-defined nuclear regions, motivating quantitative analysis of LD density and area fraction within DAPI-associated ROIs.

**Figure 11.**
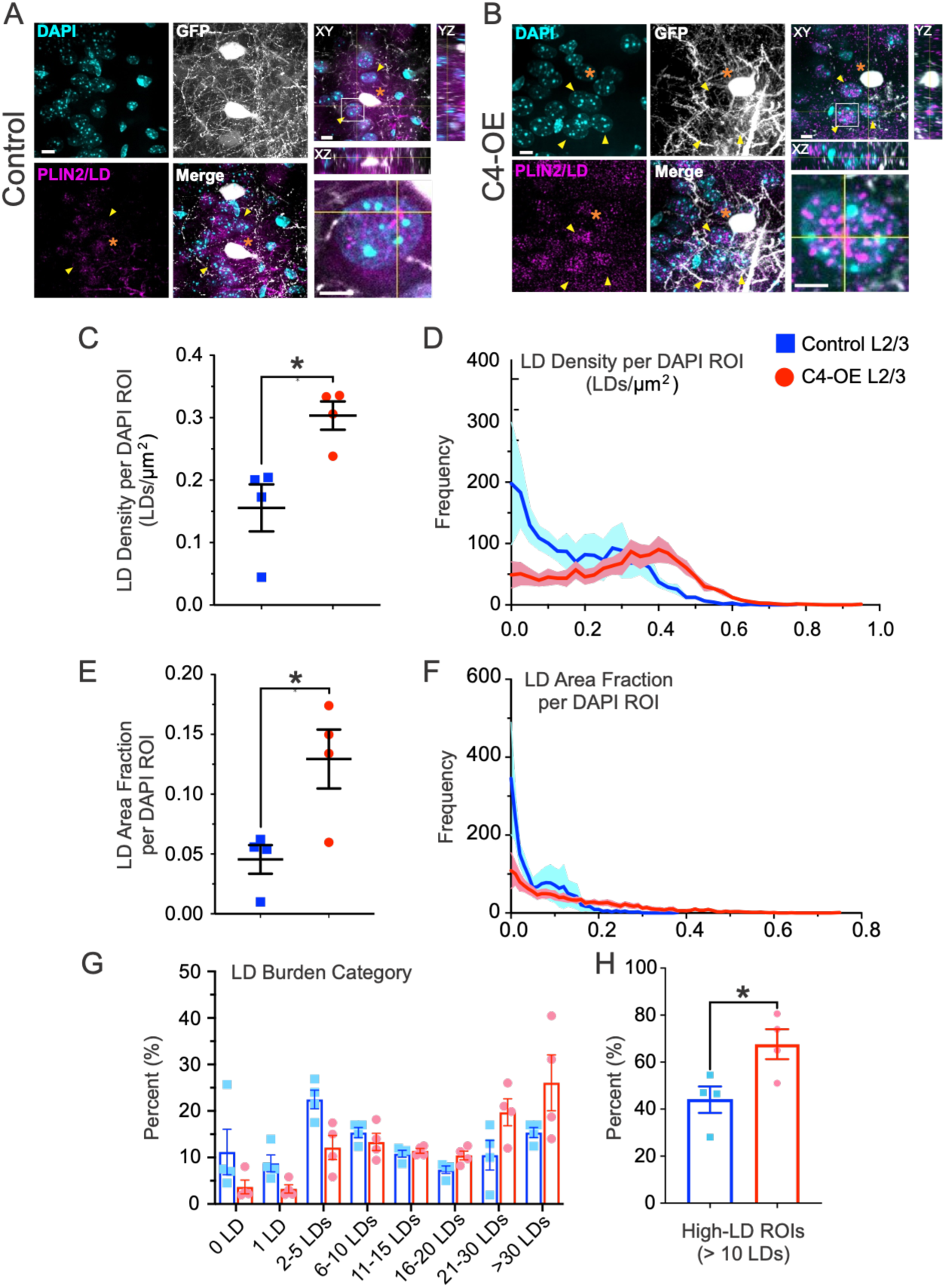
DAPI-associated perinuclear PLIN2/LD burden in L2/3 following C4-OE. (**A, B**) Representative confocal maximum-intensity projections from L2/3 regions in GFP control (**A**) and C4-OE (**B**) tissue showing DAPI, GFP, PLIN2-positive lipid droplets (PLIN2/LD), and merged channels. Channels are pseudocolored as follows: DAPI, cyan; GFP, grayscale; PLIN2/LD, magenta. Corresponding orthogonal XY, YZ, and XZ views are shown for control and C4-OE in the right panels and show the spatial relationship between PLIN2/LD puncta and DAPI-associated/perinuclear regions within the GFP-positive transfection hotspot and neighboring GFP-negative regions. Yellow arrowheads indicate PLIN2/LD puncta in GFP-negative cells, and orange asterisks indicate GFP-positive regions/cells with prominent DAPI-associated LD signal. Scale bars, 6 μm. The lower-right panels in A and B show magnified views of DAPI-associated PLIN2/LD puncta from the boxed regions in the orthogonal projections. Scale bars, 7 μm. (**C**) Quantification of LD density per DAPI ROI in GFP control (blue squares) and C4-OE (red circles) L2/3 samples. Control GFP: 0.1555 ± 0.03768 LDs/μm²; C4-OE: 0.3033 ± 0.02271 LDs/μm²; n = 4/group; Welch’s unpaired t test, P = 0.0206. (**D**) Frequency distribution of LD density per DAPI ROI. Histograms were generated from replicate-level LD density distributions using 0.025 LDs/μm² bin intervals. Values show mean frequency ± SEM across biological replicates. (**E**) Quantification of LD area fraction per DAPI ROI in GFP control and C4-OE L2/3 samples. Control GFP: 0.04545 ± 0.01197; C4-OE: 0.1294 ± 0.02464; n = 4/group; Welch’s unpaired t test, P = 0.0337. (**F**) Frequency distribution of LD area fraction per DAPI ROI. Histograms were generated from replicate-level LD area fraction distributions using 0.01 bin intervals. Values show mean frequency ± SEM across biological replicates. (**G**) Distribution of DAPI-associated ROIs across LD burden categories. Each DAPI ROI was assigned to a category based on the number of PLIN2/LD puncta detected within that ROI: 0 LDs, 1 LD, 2–5 LDs, 6–10 LDs, 11–15 LDs, 16–20 LDs, 21–30 LDs, or >30 LDs. (**H**) Quantification of high-LD ROIs, defined as DAPI ROIs containing >10 PLIN2/LD puncta. High-LD ROIs were calculated by summing the 11–15, 16–20, 21– 30, and >30 LD categories for each biological replicate. Control GFP: 44.02 ± 5.63%; C4-OE: 67.63 ± 6.38%; n = 4/group; Welch’s unpaired t test, P = 0.0327. Data are shown as mean ± SEM with individual biological replicates overlaid. LD burden was quantified in GFP-negative DAPI-associated ROIs located within the GFP-positive L2/3 transfection hotspot, representing non-transfected neighboring cells surrounding the sparsely transfected population. Measurements are interpreted as DAPI-associated/perinuclear PLIN2-positive LD burden rather than whole-cell LD content. P < 0.05. For all quantifications, DAPI ROI-level measurements were summarized within each animal, and statistical analyses were performed on animal-level summary values; individual ROIs were not treated as independent biological replicates.

To quantify this observation, we measured PLIN2-positive LD puncta density in GFP-negative DAPI-associated ROIs within the GFP-positive L2/3 transfection hotspot, thereby focusing on non-transfected neighboring cells surrounding the sparsely transfected population. C4-OE significantly increased LD density per DAPI-associated ROI by approximately 1.95-fold, corresponding to a 95% increase relative to GFP controls (**Fig. 11C**). Consistent with this increase, the LD-density distribution shifted away from the lowest-density bins toward intermediate-to-higher LD-density bins in C4-OE tissue (**Fig. 11D**).

C4-OE also significantly increased PLIN2-positive LD area fraction per DAPI-associated ROI (**Fig. 11E**). Relative to GFP controls, C4-OE produced an approximately 2.85-fold, or 185%, increase in LD area fraction. Thus, C4-OE was associated not only with increased numbers of PLIN2-positive puncta but also with a greater fraction of each analyzed DAPI-associated ROI occupied by PLIN2-positive signal. Consistent with this effect, the LD area-fraction distribution shifted away from the lowest area-fraction bins toward intermediate-to-higher values in C4-OE tissue (**Fig. 11F**).

To determine whether this increase reflected a broader shift in LD burden across DAPI-associated ROIs, we next categorized each ROI according to the number of PLIN2-positive LD puncta detected within it. Compared with GFP controls, C4-OE samples showed reduced representation in the lowest LD-burden categories and greater representation in higher-burden categories, particularly the 21–30 LD and >30 LD bins (**Fig. 11G**). We then collapsed these categories into a high-LD endpoint, defined as the percentage of DAPI-associated ROIs containing >10 PLIN2-positive LD puncta. C4-OE significantly increased the fraction of high-LD ROIs by approximately 1.54-fold, corresponding to a ∼54% increase relative to GFP controls (**Fig. 11H**). This indicates that C4-OE shifts the local ROI population toward higher-LD-burden states rather than producing only a uniform low-level increase in PLIN2 signal.

Therefore, these analyses indicate that sparse neuronal C4-OE increases perinuclear PLIN2-positive lipid-droplet burden in neighboring L2/3 cells, reflected by increases in LD density, LD area fraction, and the fraction of ROIs in high-LD-burden categories. Because the quantitative analysis focused on GFP-negative DAPI-associated ROIs within the transfection field, these findings support a non-cell-autonomous component to the lipid-associated response. In this context, increased PLIN2-positive LD burden provides an imaging-based correlate of the lipid- and metabolic-transcriptional changes observed by bulk RNA-seq and suggests that focal neuronal C4 elevation is associated with cellular remodeling extending beyond the directly transfected neurons.

## Discussion

### Sparse C4-OE Is Associated with Detectable Local Transcriptomic Remodeling

Despite sparse neuronal transfection in our IUE model, bulk RNA-seq revealed robust transcriptional changes in microdissected cortical tissue, including strong upregulation of C4 itself. At first glance, this may seem counterintuitive given the dilution inherent to bulk profiling. However, several factors likely contribute to this signal. First, because endogenous C4 expression is normally low in this context (9,17,18), even a small population of C4-overexpressing cells can generate a substantial tissue-level fold-change. Second, focal neuronal C4 elevation may be associated with transcriptional responses extending beyond the directly transfected neurons, including programs associated with neighboring neuronal and non-neuronal populations. Third, C4-associated alterations in neuronal activity, synaptic organization, or homeostatic signaling may contribute to broader changes in local gene-expression programs (7,71).

Together, these considerations help explain how a spatially restricted manipulation can be associated with robust changes in bulk transcriptomic profiles. Rather than viewing the bulk RNA-seq signal solely as a diluted readout of the sparsely transfected population, the data support a working model (**S4 Fig.**) in which focal neuronal C4 elevation is associated with broader transcriptional and metabolic changes within the local cortical field. This interpretation is further supported by the reference-based projection, MFISH validation, and PLIN2/lipid-droplet analyses, which together indicate that sparse neuronal C4-OE is associated with changes extending beyond the directly targeted neuronal population.

### C4-OE Is Associated with Cholesterol Biosynthesis and Broader Homeostatic Transcriptional Programs

One of the strongest and most consistent transcriptional features associated with sparse neuronal C4 elevation was the positive enrichment of cholesterol- and sterol-biosynthesis-associated programs. This signal was evident at the level of individual DEGs, pathway enrichment, and co-expression analysis, with coordinated upregulation of genes spanning multiple steps of the cholesterol biosynthetic pathway. This finding is particularly relevant in the brain, where cholesterol is synthesized largely within the CNS because peripheral cholesterol has limited access across the blood–brain barrier (57,72,73). Cholesterol is required not only for myelin production but also for neuronal membrane organization, synapse formation and maintenance, synaptic vesicle release, and neurotransmission (66,74,75). This is especially relevant given our previous findings that neuronal C4 elevation alters excitatory connectivity and AMPA-receptor trafficking, processes that depend on membrane composition and lipid homeostasis.

The cholesterol-associated response was notable for its consistency across analytical approaches. Ten of the fifteen genes represented in the WikiPathways cholesterol biosynthesis set were significantly upregulated, including *Hmgcs1, Hmgcr, Cyp51, Idi1, Msmo1, Mvd, Sqle, Dhcr7, Sc5d,* and *Nsdhl*, and cholesterol/sterol biosynthesis was among the strongest positively enriched programs by GSEA and GO analysis. These genes span the mevalonate and downstream sterol biosynthetic pathways, which provide cholesterol as well as intermediate metabolites with broader roles in membrane biology and cellular signaling (76,77). A cholesterol-associated WGCNA module at the reference power of 15 also showed a strong genotype association and enrichment for cholesterol biosynthetic processes. Because gene-level module assignments varied across alternative network solutions, we interpret individual WGCNA module membership cautiously and view the network analysis as supportive rather than independently confirmatory evidence. Thus, the convergence of differential-expression, pathway-enrichment, and co-expression analyses supports cholesterol biosynthesis as one of the more consistent transcriptional features associated with sparse neuronal C4 elevation.

At the transcriptional level, this response appeared preferentially centered on cholesterol biosynthesis rather than a uniform shift across all aspects of cholesterol homeostasis. We did not detect comparable transcriptional regulation of pathways associated with cholesterol export, neuronal uptake, oligodendrocyte cholesterol processing or myelination, or neuronal cholesterol clearance (78–84). These processes could still be influenced at post-transcriptional, post-translational, or metabolic levels, but they were not detectably regulated at the transcript level in the current bulk RNA-seq dataset.

The lipid-associated response was not limited to cholesterol biosynthesis. C4-OE was also associated with altered expression of lipid-droplet and lipid-turnover genes, including downregulation of *Plin4*, *Plin3*, and other lipid-metabolism-associated transcripts, together with negative enrichment of fatty-acid metabolism and mitochondrial fatty-acid β-oxidation programs. Lipid droplets are dynamic organelles involved in lipid storage and buffering, cellular stress responses, and immune-metabolic regulation rather than simply passive lipid depots (58,67,69). Recent work indicates that lipid-droplet abundance alone does not distinguish protective from detrimental states, as APOE2- and APOE4-associated droplets can accumulate similarly yet differ in proteomic composition and turnover; future studies should therefore examine the molecular contents and dynamics of C4-associated lipid droplets (85). These transcriptional findings therefore suggest that sparse neuronal C4 elevation is associated with broader changes in lipid handling in addition to increased cholesterol biosynthesis.

The broader transcriptional response also included positively enriched synaptic organization, axon-related, vascular/endothelial, adhesion, extracellular-matrix, and interferon/inflammatory-associated programs, whereas synaptic translation, ribosomal, and fatty-acid oxidation-related programs were negatively enriched. The opposing regulation of synaptic organization and synaptic translation suggests that C4 elevation is associated with changes in multiple aspects of neuronal homeostasis rather than a uniform increase or decrease in synaptic gene expression. This interpretation is consistent with established roles of activity-dependent and homeostatic mechanisms in adapting synaptic function to altered circuit activity (71), as well as with altered expression of genes such as *Homer1* and *Rheb*, which participate in synaptic organization, axonal growth, and mTOR-associated signaling (59–62). Likewise, enrichment of cell-cycle-, EMT-, and MYC-related gene sets in the predominantly post-mitotic P21 cortex should be interpreted as transcriptional signatures that may reflect non-neuronal-, vascular-, immune/stress-, or tissue-remodeling-associated programs rather than neuronal proliferation.

The immune-associated response was heterogeneous rather than uniformly directional. Although all three genes encoding the C1q complex (*C1qa, C1qb,* and *C1qc*) and several other immune-associated transcripts were reduced, interferon-, cytokine-, and inflammatory-response gene sets were positively enriched. Thus, the transcriptional response to sparse C4 elevation is better interpreted as involving directionally distinct immune- and stress-associated programs rather than generalized activation or suppression of inflammation. Importantly, the functional consequences of these transcriptional changes at the protein and pathway-activity levels remain to be determined. This distinction is particularly relevant for the complement system, in which transcript abundance of individual components may not directly reflect protein abundance, proteolytic activation, complex formation, or overall pathway activity.

Recent work showing that glial cholesterol supply can influence neuronal synaptic gene-expression programs, together with reciprocal transcriptional responses in glia, provides a precedent for lipid-homeostatic changes being coupled across cell populations rather than confined to a single cellular compartment (86). Because C4 elevation was restricted to a sparse neuronal subset, the broader lipid-associated transcriptional response observed here is consistent with changes extending beyond the directly targeted neurons, although the bulk RNA-seq data alone do not identify the cellular sources of these signals.

Because these experiments were performed in outbred CD-1 IGS mice, strain-specific genetic variation could influence individual transcript expression. However, there is currently limited evidence that the major pathway-level signatures emphasized here—particularly cholesterol biosynthesis, synaptic organization, vascular/ECM, and immune/stress-associated programs— would be expected to differ systematically from those observed in commonly used inbred strains. Moreover, across our prior studies using the same sparse neuronal C4-OE framework, Comer et al. (9) and Phadke et al. (17) identified reproducible molecular, cellular, circuit, and behavioral phenotypes without evidence of unusually high variability across these readouts. We therefore interpret isolated low-count or potentially polymorphic loci cautiously and place greater emphasis on convergent pathway-level findings and independently validated phenotypes.

Sex composition is also a limitation of the current RNA-seq design. Although the study was not prospectively designed or powered to test sex-dependent effects, sex was included as an additive covariate in the differential-expression model, and the major C4-associated pathway-level findings remained evident. Several key results were also consistent across individual biological replicates rather than being driven by a single animal: C4-OE and control samples separated primarily along PC1, cholesterol-associated genes such as *Hmgcr* and *Hmgcs1* showed consistent expression differences across samples, and the cholesterol-associated WGCNA eigengene displayed opposing expression patterns in all four controls versus all four C4-OE animals. Together with our prior sparse neuronal C4-OE studies (9,17), in which we observed molecular, cellular, synaptic, circuit, and behavioral phenotypes without clear sex-dependent differences, these findings make sex composition unlikely to be a sufficient explanation for the overall C4-associated phenotype. Nevertheless, the present design cannot resolve sex-by-genotype interactions or exclude the possibility that sex modulates the magnitude or composition of specific transcriptional responses. Appropriately powered, sex-balanced cohorts will be required to address that question directly.

Overall, these findings identify cholesterol biosynthesis as a particularly consistent component of the C4-OE transcriptional response, embedded within a broader pattern of lipid, synaptic, vascular/ECM, and immune/stress-associated changes. This broader organization suggests that sparse neuronal C4 elevation is associated with coordinated lipid-, synaptic-, vascular-, and immune/stress-related transcriptional changes rather than a single inflammatory or synaptic response.

### Reference Projection and Spatial Validation Identify Cell-State- and Lipid-Associated Responses to Sparse Neuronal C4 Elevation

The reference-based projection provides additional context for the broad transcriptional response detected by bulk RNA-seq. The directional C4-OE signatures mapped onto distinct cortical reference spaces: downregulated genes were prominently associated with an astrocyte-associated PC1 axis, whereas upregulated genes mapped onto a vascular-associated PC1 axis characterized by mural- and endothelial-associated structure. PC2 retained additional neuronal-associated organization, including prominent L2/3 IT-associated structure in the downregulated projection. This latter feature is particularly relevant because L2/3 IT neurons are the reference population most closely corresponding to the upper-layer excitatory neurons targeted by IUE. These patterns indicate that the C4-OE transcriptional response contains both neuronal-associated structure related to the manipulated population and broader non-neuronal-associated signatures.

The correlation-domain analysis further refined this organization. The downregulated signature contained an astrocyte-associated lipid–metabolic stress domain and a complement/immune-stress-response domain, whereas the upregulated signature contained vascular/endothelial–interferon-associated, C4b-containing extracellular matrix-associated, and cholesterol/lipid biosynthesis domains. These patterns are consistent with the broader pathway-level analyses and suggest that the directional C4-OE response maps onto multiple cortical cell-state-associated programs rather than a single neuronal or inflammatory transcriptional response. Recent work in an α-synuclein model similarly points to C4-dependent neuron–astrocyte crosstalk, with neuronal C4 induction enhanced by astrocytes and C4 exposure amplifying astrocytic inflammatory responses (87). Importantly, these projections do not establish the cellular origin of the bulk RNA-seq changes or indicate altered cell-type abundance; they identify transcriptional correspondence with defined cortical reference populations.

The astrocyte-associated component is particularly relevant to the lipid-homeostatic findings because astrocytes are major regulators of brain cholesterol synthesis, lipid handling, metabolic support, extracellular homeostasis, and synaptic function (88,89). Recent work further suggests that cholesterol availability can influence reciprocal transcriptional programs between glia and neurons (86). However, the projection alone does not demonstrate that astrocytes are the source of the altered transcripts. Rather, it provides a framework for interpreting the spatial validation experiments, which directly tested whether selected molecular changes were detectable in cells neighboring the manipulated neuronal population.

MFISH provided such spatial support. Although *Hmgcr* and *Fam107a* did not differ significantly between GFP-positive C4-OE and GFP-control neurons, neighboring GFP-negative cells within the C4-OE field showed significantly increased *Hmgcr* expression and a strong trend toward reduced *Fam107a*. Thus, for the genes examined, the clearest *C4*-associated expression differences were detected in neighboring non-transfected cells rather than in the directly manipulated neurons. This finding provides single-cell spatial support for the broader bulk RNA-seq response and supports a C4-OE-associated non-cell-autonomous component of local transcriptional remodeling.

The PLIN2 imaging experiments provided an independent cellular correlate of this lipid-associated response. Within GFP-negative DAPI-associated regions in the local C4-OE field, PLIN2-positive lipid-droplet density, area fraction, and the proportion of high-lipid-droplet-burden regions were all increased. Lipid droplets are dynamic organelles involved in lipid storage, buffering, cellular stress responses, and immune-metabolic regulation (67,69,85). Notably, the PLIN2 imaging phenotype represents an independent protein- and organelle-associated readout and should not be expected to directly mirror *Plin2* transcript abundance, as lipid-droplet-associated proteins can be influenced by post-transcriptional regulation, protein stability and turnover, and compensatory cellular responses. Importantly, lipid-droplet abundance alone does not distinguish adaptive from detrimental states, as protective APOE2- and risk-associated APOE4 astrocytes can both accumulate lipid droplets while differing in lipid-droplet proteomic composition and turnover (85). Future studies should therefore examine the molecular composition, proteomic state, and turnover dynamics of the C4-associated lipid droplets.

The cellular identity underlying the PLIN2 phenotype remains unresolved because the current analysis used DAPI-associated/perinuclear regions rather than cell-boundary or cell-type-specific markers. The increased lipid-droplet burden could therefore arise from astrocytes, microglia, neurons, vascular-associated cells, or multiple neighboring populations. Likewise, the vascular-associated projection should be interpreted as correspondence with mural- and endothelial-associated transcriptional programs rather than direct evidence of altered vascular-cell abundance or state. Future single-nucleus RNA-seq, spatial transcriptomics, and multiplex imaging incorporating neuronal, astrocytic, microglial, and vascular markers will be needed to identify the cellular states carrying these responses.

Taken together, the convergence of reference-based projection, MFISH, and PLIN2 imaging supports a model in which sparse neuronal C4 elevation is associated with non-cell-autonomous remodeling of the local cortical field, with transcriptional and lipid-associated changes extending beyond the directly targeted neurons. These findings broaden the consequences associated with neuronal C4 elevation to the surrounding cortical environment, while leaving the precise cellular sources, mechanisms, and functional consequences to be resolved.

### Sparse C4-OE Shows That a Single Immune-Risk Gene Can Influence the Local Cortical Environment

A key implication of this study is that sparse neuronal C4 elevation is associated with broad molecular and cellular changes in the absence of broader disease pathology. In Alzheimer’s disease, SCZ, aging, and other complex disease contexts, complement dysregulation occurs alongside inflammation, neuronal dysfunction, vascular changes, altered activity, and progressive circuit pathology (90,91), making it difficult to distinguish whether complement changes are primary contributors, compensatory adaptations, or secondary consequences of disease. By contrast, our sparse C4-OE model introduces a defined immune-associated perturbation into an otherwise intact developing cortical circuit. The resulting cholesterol-biosynthesis signature, synaptic- and vascular-associated transcriptional programs, astrocyte-associated reference structure, neighboring-cell *Hmgcr* induction, and increased PLIN2-positive lipid-droplet burden support the idea that focal neuronal C4 elevation is associated with effects extending across the local cortical field.

Complement is well positioned to generate such local field effects because its activity is spatially restricted, amplified, and regulated (19,91). In peripheral tissues, complement can shape responses around altered or damaged targets while regulatory mechanisms limit inappropriate spread to neighboring healthy cells (19). Our findings raise the possibility that an analogous spatial principle operates in the developing cortex, where sparse neuronal C4 elevation is associated with transcriptional and lipid-related changes in surrounding cells. Together with previous studies showing that neuronal C4 elevation alters synaptic connectivity, receptor trafficking, circuit function, and SCZ-relevant behavior (9,10,17,18), these findings broaden the interpretation of C4 biology beyond synapse elimination alone and support a model in which focal C4 dysregulation contributes to a wider local cortical state relevant to circuit maturation and vulnerability.

## Acknowledgments

We thank members of the Cruz-Martín lab and Dr. Borislav Dejanovic and Dr. Sarah Melzer for their critical reading of the manuscript and helpful discussions, as well as Dr. Ashley L. Comer for performing electroporations and Dr. Tushare Jinadasa for conducting tissue microdissection. Imaging was performed in the Advanced Light Microscopy Core facility of the NeuroTechnology Center at the University of Colorado Anschutz Medical Campus.

## Use of AI-Assisted Tools

During manuscript preparation, AI-assisted tools, including OpenAI ChatGPT, Anthropic Claude, and Google Gemini, were used to support language editing, manuscript section organization, author contribution language, figure legend refinement, and clarity of phrasing. These tools were also used to help improve clarity and consistency in the description of computational workflows and results. AI-assisted tools were not used to generate primary data, perform final statistical analyses, create original scientific conclusions, or replace author interpretation. No AI-assisted tool is listed as an author. All AI-assisted text and suggestions were reviewed, edited, and approved by the authors, who take full responsibility for the accuracy, integrity, and final content of the manuscript.

## Author Contribution Statement

M.S.C., S.B., and J.Á.P. contributed equally as co-first authors. M.S.C. provided substantial conceptual input throughout the study, performed imaging experiments and data analysis, led key spatial validation experiments including MFISH and PLIN2/LD imaging analyses, contributed to data interpretation and figure development, and helped shape and revise the manuscript. S.B. designed the initial computational analysis pipeline, coordinated major aspects of the study, performed the majority of the bulk RNA-seq and downstream transcriptomic analyses, generated figures, and wrote the initial draft of the manuscript. J.Á.P. provided substantial conceptual input, contributed to experimental design and interpretation of the C4-OE model, performed imaging experiments and data analysis, helped integrate the transcriptomic, spatial, and lipid-remodeling findings, contributed to figure development, and participated in manuscript writing and revision. Y.A.S. and C.G.-D. performed the cell-type-aware projection and correlation-domain analyses, including writing the associated Methods and Results sections and generating the corresponding figures in collaboration with A.C.-M. D.G.S. and S.V.C. contributed to imaging, image analysis, and validation experiments. S.E.C.S. optimized the analysis pipeline, reanalyzed portions of the dataset, contributed to figure generation, and helped refine the manuscript. R.A.P. performed the microdissections, helped curate the initial data, organized the shared repository for collaborative access, and contributed to figure development and manuscript editing. A.B. supported data curation and repository organization. A.H.-K. optimized the PLIN2/LD staining protocol and imaging parameters and contributed to data interpretation. P.C. contributed to manuscript editing and revision. C.D. provided conceptual input and helped interpret the findings. A.C.-M. conceived and supervised the project, secured funding, directed the overall study design, and co-wrote the manuscript with M.S.C., S.B., J.Á.P., and other authors, incorporating feedback from all coauthors. All authors reviewed and approved the final manuscript.

## Data Availability Statement

Analysis code, the raw gene-level RNA-seq count matrix, and the processed count matrix are available at https://github.com/Cruz-Martin-Lab/C4-OE-CorticalRNAseq. Third-party datasets used in the analyses, including the Aryal et al. schizophrenia synaptic proteome, the Allen Brain Map SMART-seq cortical reference, and MSigDB mouse collections v2023.2, are available from their original sources. MFISH image quantification was performed using the previously published FijiFISH-based workflow described by Sullivan et al. (2025). Custom Fiji macros used for the PLIN2/lipid-droplet image analysis are available from the corresponding author upon reasonable request.

## Ethics, Consent to Participate, and Consent to Publish

Declarations: not applicable.

## Supporting Information

### Supplementary Figure Legends

**S1 Fig.**
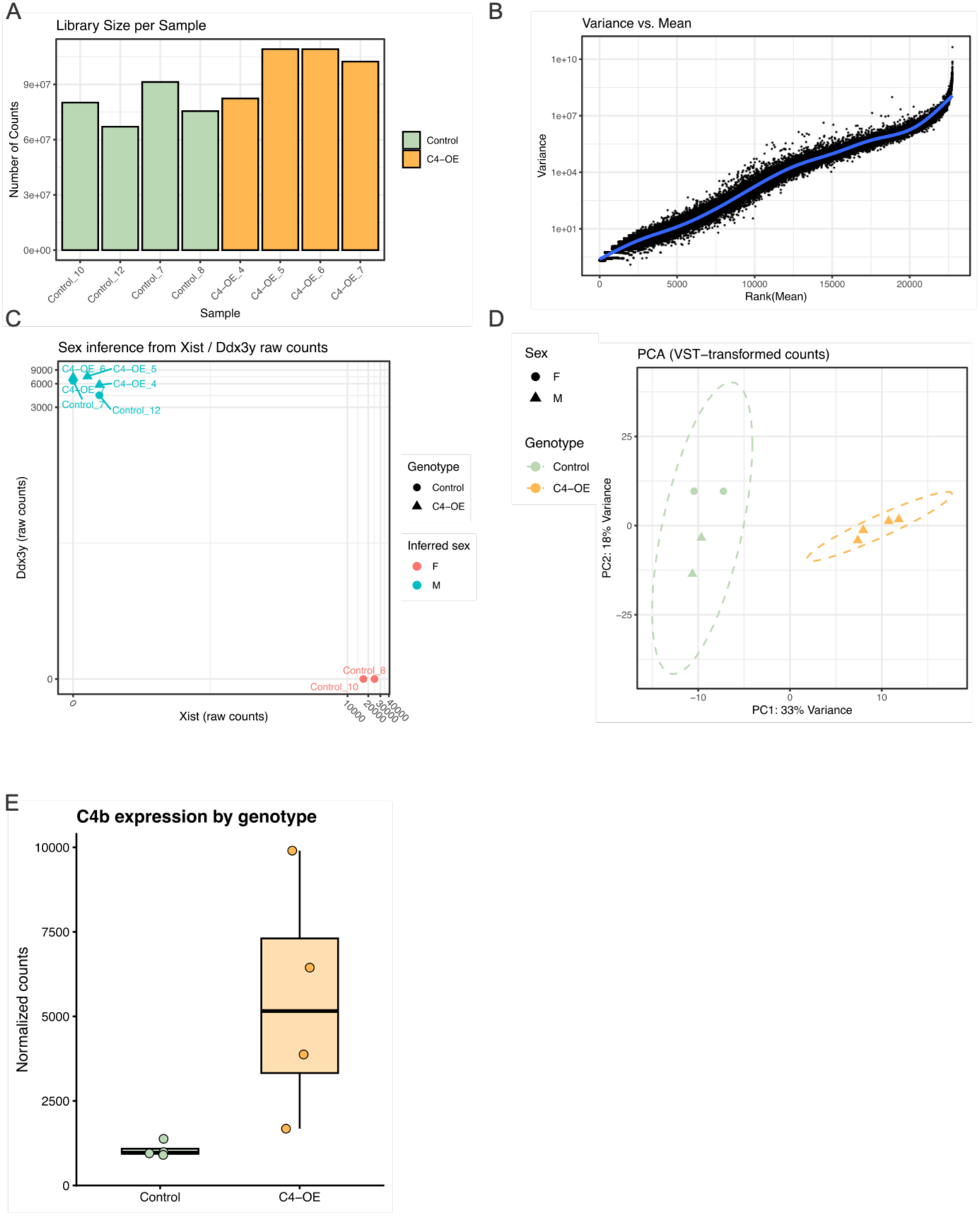
RNA-seq quality control, sample characteristics, and *C4b* expression in Control and C4-OE mPFC samples. (**A**) Library size for each RNA-seq sample, shown as the total number of sequencing counts. Bars are colored by genotype (Control, pale green; C4-OE, orange). Four biological replicates were analyzed per group. (**B**) Gene-level variance plotted against the rank of mean expression across samples, illustrating the mean–variance relationship in the RNA-seq dataset. The blue line represents the smoothing curve generated by ggplot2::geom_smooth() and is included as a visual guide to the overall trend; it does not represent the DESeq2 dispersion model. (**C**) Sex inference based on raw counts of *Xist* and the Y-linked transcript *Ddx3y*. Samples with high *Xist* and low *Ddx3y* expression were inferred as female, whereas samples with low *Xist* and high *Ddx3y* expression were inferred as male. Control samples comprised two females and two males, whereas all four C4-OE samples were male. Genotype is indicated by point shape and inferred sex by color. (**D**) Principal component analysis of variance-stabilized expression values. PC1 and PC2 explained 33% and 18% of the variance, respectively. Samples are colored by genotype (Control, pale green; C4-OE, orange), and point shape indicates sex (female, circles; male, triangles). Dashed ellipses delineate samples by genotype. (**E**) DESeq2-normalized C4b counts in Control and C4-OE samples, with individual biological replicates overlaid. *C4b* expression was significantly increased in C4-OE tissue (log₂FC = 2.53; adjusted P = 4.93 × 10⁻³), consistent with successful detection of the intended experimental perturbation.

**S2 Fig.**
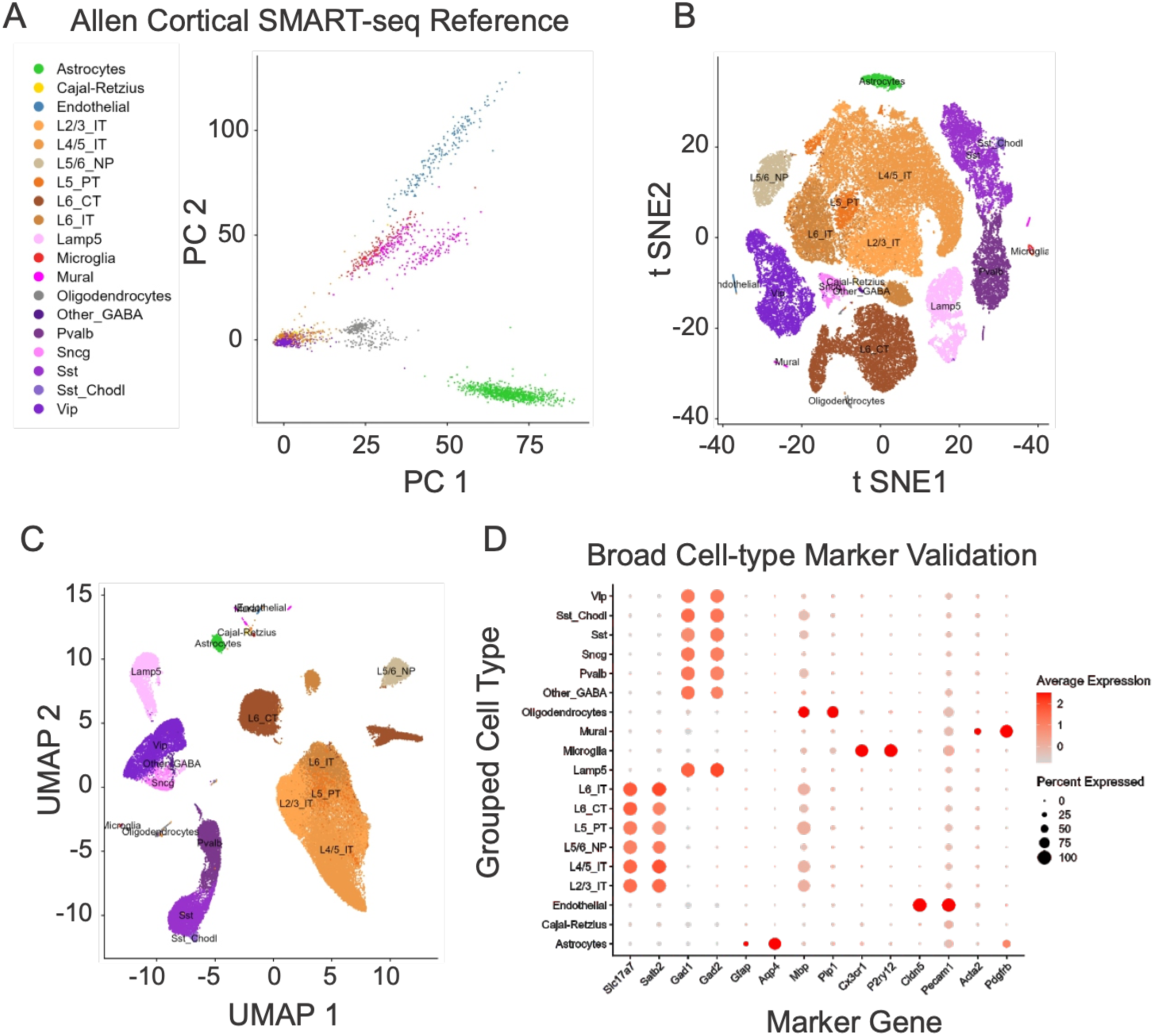
Construction and validation of the curated cortical single-cell reference dataset. (**A**) Principal component analysis (PCA) of the curated Allen Mouse Whole Cortex and Hippocampus SMART-seq reference dataset after removal of hippocampal neuronal populations and consolidation of cortical cell classes. Each point represents a single reference cell colored by grouped cortical cell identity. (**B**) t-SNE visualization of the same curated reference dataset showing separation of major neuronal and non-neuronal cortical cell classes. (**C**) UMAP visualization of the curated cortical reference, demonstrating clustering of excitatory neuron subclasses, inhibitory interneuron subclasses, and non-neuronal populations. (**D**) Dot plot showing expression of broad cell-type marker genes across grouped cortical cell classes, used to confirm the expected identity of major reference populations. Dot size represents the percentage of cells expressing each marker, and color intensity represents average scaled expression. Cell classes include astrocytes, Cajal-Retzius cells, endothelial cells, microglia, mural/perivascular cells, oligodendrocytes, excitatory neuron subclasses, and inhibitory interneuron subclasses. Abbreviations: L2/3_IT, layer 2/3 intratelencephalic excitatory neurons; L4/5_IT, layer 4/5 intratelencephalic excitatory neurons; L5/6_NP, layer 5/6 near-projecting excitatory neurons; L5_PT, layer 5 pyramidal tract excitatory neurons; L6_CT, layer 6 corticothalamic excitatory neurons; L6_IT, layer 6 intratelencephalic excitatory neurons; Lamp5, lysosomal-associated membrane protein family member 5 interneurons; Other_GABA, Meis2-positive/other GABAergic interneurons; Pvalb, parvalbumin interneurons; Sncg, gamma-synuclein interneurons; Sst, somatostatin interneurons; Sst_Chodl, somatostatin/chondrolectin interneurons; Vip, vasoactive intestinal peptide interneurons. This curated reference was used for cell-type-aware projection of C4-OE-regulated bulk RNA-seq signatures.

**S3 Fig.**
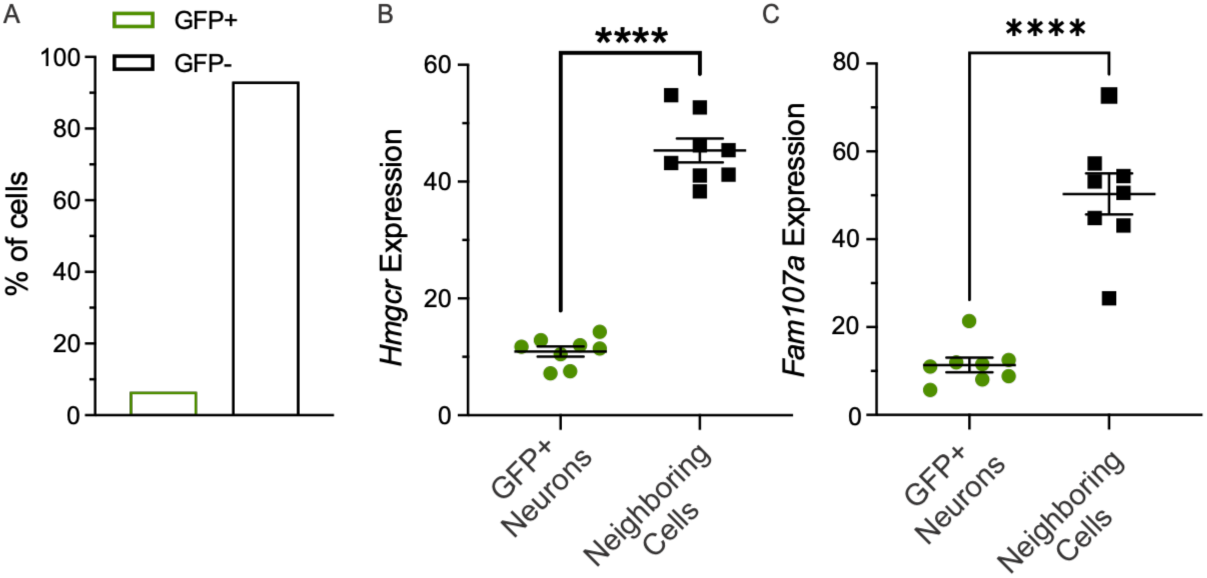
Electroporation efficiency and pooled comparison of GFP-positive and GFP-negative cells. (**A**) Proportion of GFP-positive and GFP-negative cells across sampled tissue sections quantified by MFISH. GFP-positive cells represented 6.67% of the total sampled cell population within the local transfection hotspot analyzed here, whereas GFP-negative cells represented 93.33%. (**B**) Quantification of *Hmgcr* expression, measured as percent area covered, in GFP-positive neurons and neighboring GFP-negative cells pooled across GFP control and C4-OE conditions. GFP-positive neurons: 10.94 ± 0.87%; neighboring GFP-negative cells: 45.36 ± 2.04%. Data are shown as mean ± SEM. Welch’s unpaired t test, ****P < 0.0001. (**C**) Quantification of *Fam107a* expression, measured as percent area covered, in GFP-positive neurons and neighboring GFP-negative cells pooled across GFP control and C4-OE conditions. GFP-positive neurons: 11.38 ± 1.65%; neighboring GFP-negative cells: 50.33 ± 4.67%. Data are shown as mean ± SEM. Welch’s unpaired t test, ****P < 0.0001.

**S4 Fig.**
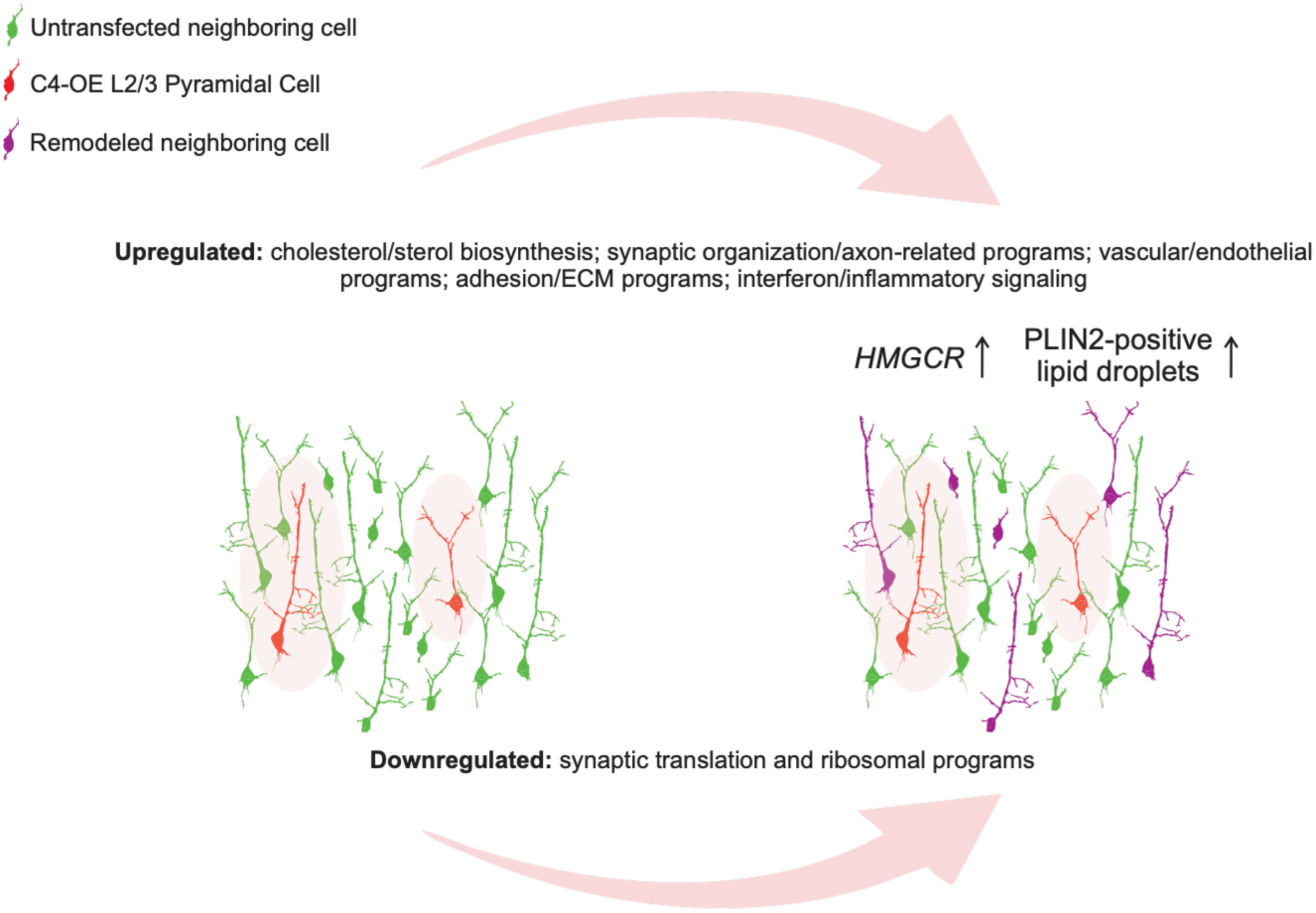
Working model of sparse neuronal C4 elevation and remodeling of the local cortical microenvironment. Model summarizing the transcriptional and cellular responses associated with sparse neuronal C4 elevation in a small subset of layer 2/3 pyramidal neurons. Sparse neuronal C4 elevation is associated with coordinated changes across the surrounding cortical tissue, including increased cholesterol and sterol biosynthesis, positive enrichment of synaptic organization and axon-related programs, increased vascular/endothelial, adhesion/extracellular-matrix, and interferon/inflammatory-associated transcriptional programs, and downregulation of synaptic translation and ribosomal programs. The model also incorporates increased *Hmgcr* expression and increased PLIN2-positive lipid-droplet burden in neighboring GFP-negative cells within the C4-OE cortical field. Together, these findings support a model in which sparse neuronal C4 elevation is associated with broad remodeling of molecular programs extending beyond the directly manipulated neuronal population.

**Striking Image.**
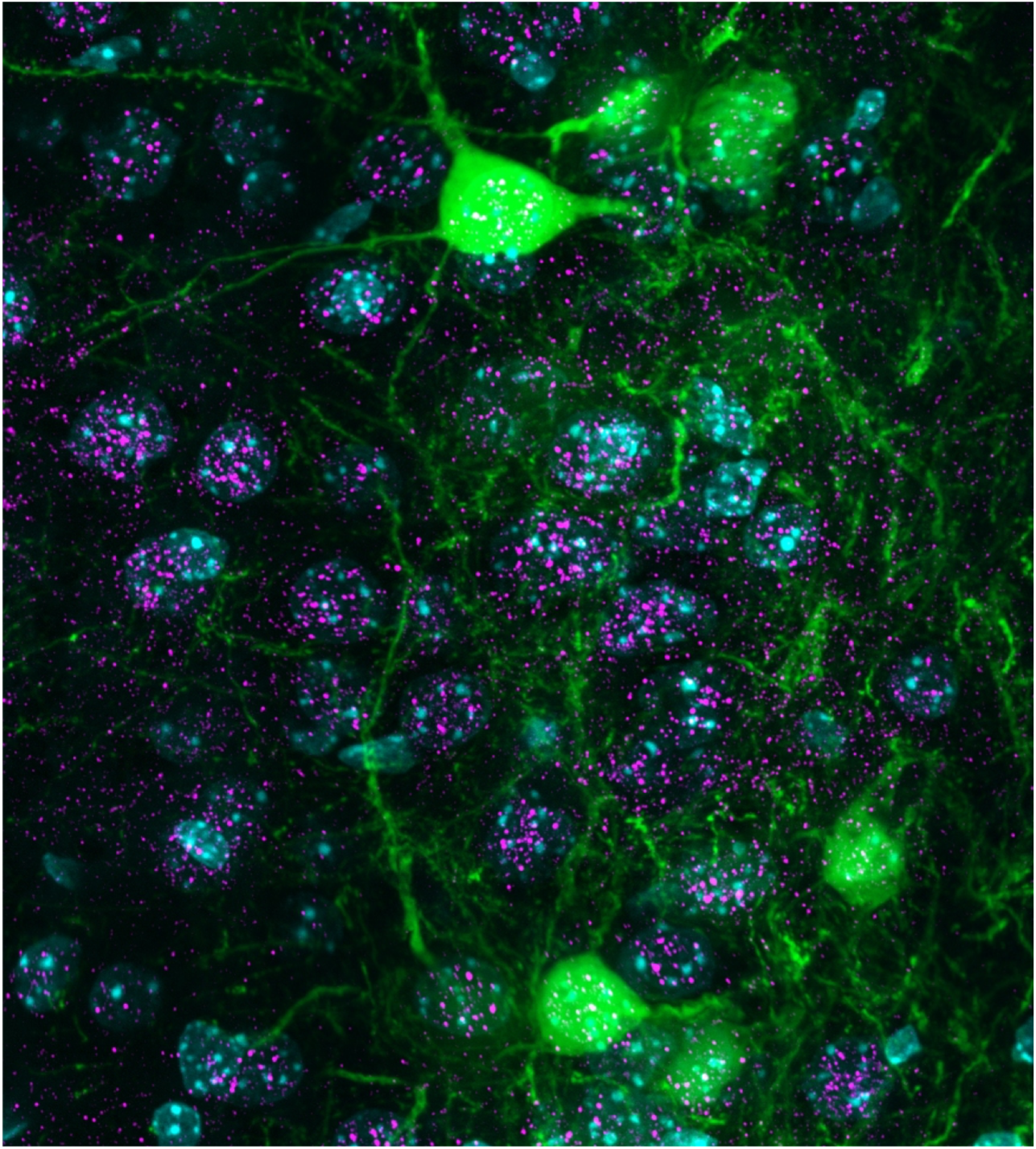
Sparse neuronal C4 elevation and PLIN2-positive lipid droplets in the cortical field. Representative fluorescence image showing GFP-positive layer 2/3 pyramidal neurons and their processes within the transfected cortical region. GFP is shown in green, DAPI-labeled nuclei in cyan, and PLIN2-positive lipid droplets in magenta. PLIN2-positive lipid droplets are visible throughout the local transfection field, including in association with GFP-positive regions and surrounding GFP-negative neighboring cells, illustrating that lipid-droplet accumulation extends beyond the directly transfected neuronal population**. Image credit:** Juan E. Ávila-Pagán and Alberto Cruz-Martín, Cruz-Martín laboratory, Department of Anesthesiology, University of Colorado Anschutz Medical Campus.

